# Reconfiguration of the visual code and retinal cell type complement in closely related diurnal and nocturnal mice

**DOI:** 10.1101/2024.06.14.598659

**Authors:** Annette E Allen, Joshua Hahn, Rose Richardson, Andreea Pantiru, Josh Mouland, Beatriz Baño-Otalora, Aboozar Monavarfeshani, Wenjun Yan, Christopher Williams, Jonathan Wynne, Jessica Rodgers, Nina Milosavljevic, Patrycja Orlowska-Feuer, Riccardo Storchi, Joshua R. Sanes, Karthik Shekhar, Robert J Lucas

## Abstract

How does evolution act on neuronal populations to match computational characteristics to functional demands? We address this problem by comparing visual code and retinal cell composition in closely related murid species with different behaviours. *Rhabdomys pumilio* are diurnal and have substantially thicker inner retina and larger visual thalamus than nocturnal *Mus musculus*. High-density electrophysiological recordings of visual response features in the dorsal lateral geniculate nucleus (dLGN) reveals that *Rhabdomys* attains higher spatiotemporal acuity both by denser coverage of the visual scene and a selective expansion of elements of the code characterised by non-linear spatiotemporal summation. Comparative analysis of single cell transcriptomic cell atlases reveals that realignment of the visual code is associated with increased relative abundance of bipolar and ganglion cell types supporting OFF and ON-OFF responses. These findings demonstrate how changes in retinal cell complement can reconfigure the coding of visual information to match changes in visual needs.

## Introduction

Vertebrate central nervous systems share common structural, developmental and genetic blueprints. This standard architecture must accommodate species-specific differences in dominant sense(s), modes of locomotion, and strategies for food seeking, predator avoidance, and reproduction. Such profound differences in functional requirements can exist for species that are closely related by phylogeny, highlighting the need for plasticity in the common blueprint of the brain.^1^ Recent technological developments allow exploration of how such adaptations arise with unprecedented resolution. On the one hand, large-scale recordings of neuronal populations facilitate an unbiased comparison of information channels across species^2,3^, linking coding strategies to ecological niches. On the other, high-throughput single cell transcriptomics allows comprehensive identification of cell types and cross-species comparisons^4–6^. With these techniques in hand, it becomes possible to describe the process of central nervous system evolution by tracing changes in cell type complement and adaptations in computational characteristics.

Here we apply this approach to a comparison of the early visual system in two closely related murid rodents. The laboratory mouse (*Mus musculus*) is a popular model for understanding mammalian vision in health and disease and for establishing general principles of neural function. Like most murid species, *Mus* are predominantly nocturnal, and this is reflected in key features of the visual system, including a rod dominant retina and a UV transmitting lens^7^. However, some Muridae are diurnal, including the four striped mouse, *Rhabdomys pumilio*, of sub-saharan Africa^8,9^. The switch to diurnality in *Rhabdomys* has been associated with substantial changes in visual system anatomy^10,11^. The *Rhabdomys* retina is cone-dominated, its lens absorbs UV light, and both its retina and visual centres in the brain are expanded in volume compared to *Mus*^10–12^. Moreover, an unbiased analysis of the *Rhabdomys* genome reveals that genes involved in vision exhibit accelerated evolution compared to related murids^13^.

*Rhabdomys and Mus* represent a case study in how differences in temporal niche are reflected in visual system anatomy^14^. However, an important unanswered question is how such anatomical expansion impacts the visual code. In principle, the enhanced capacity to process visual information in *Rhabdomys* could allow: the same computations to be performed with higher precision; a wider array of visual features to be extracted, describing the scene with greater granularity; or reconfigurations of the visual code towards features of particular ecological importance. Similarly, contributors to the anatomical expansion remain unclear. For example, both inner nuclear and ganglion cell layers are thicker in the *Rhabdomys* retina^10–12^, but the increased cell number might be accompanied by altered frequencies of cell types shared with *Mus,* emergence of new types, or both. Addressing these questions will help reveal how the blueprint of neural circuits can be adjusted to align computation to changing ecology.

To address these issues, we combined high-density electrophysiological recordings of visual response properties in the dorsal lateral geniculate nucleus (dLGN; primary visual thalamus) of *Mus* and *Rhabdomys* with comparative analyses of retinal cell atlases from these species. We find that the enhanced capacity of the *Rhabdomys* visual system is applied not only to sample the scene with higher density but also to facilitate a realignment towards nonlinear spatiotemporal summation. This functional realignment can be traced to adjustments in the relative abundance of multiple bipolar and retinal ganglion cell types. Our data thus show how evolution can shape computation by selective expansion of cell types within the framework of a common neural circuit blueprint.

## Results

### Realignment in thalamocortical visual code between *Mus* and *Rhabdomys*

Based on the assumption that thalamocortical vision may more reliably reflect inter-species differences in visual ecology than parts of the visual system responsible for reflexive behaviours such as optomotor coordination, pupil regulation and circadian entrainment, we set out to compare the visual code in *Rhabdomys* and *Mus* dorsal lateral geniculate nucleus (dLGN; principal relay of visual information to the cortex). Using a 256-channel recording electrode across multiple placements in both sagittal and coronal orientations we recorded visually evoked activity for neurons across the dLGN of both species (Figure 1a; Supplementary Figure 1). To broadly survey the diversity of visual responses in the two species, we applied a full field temporal modulation stimulus (Figure 1b) previously applied in both the retina and dLGN of *Mus* to identify functional response types^15^. dLGN neurons in both species showed strong yet diverse responses to this stimulus (n=611 and n=422 units in *Rhabdomys* and *Mus,* respectively; Figure 1).

**Figure 1.**
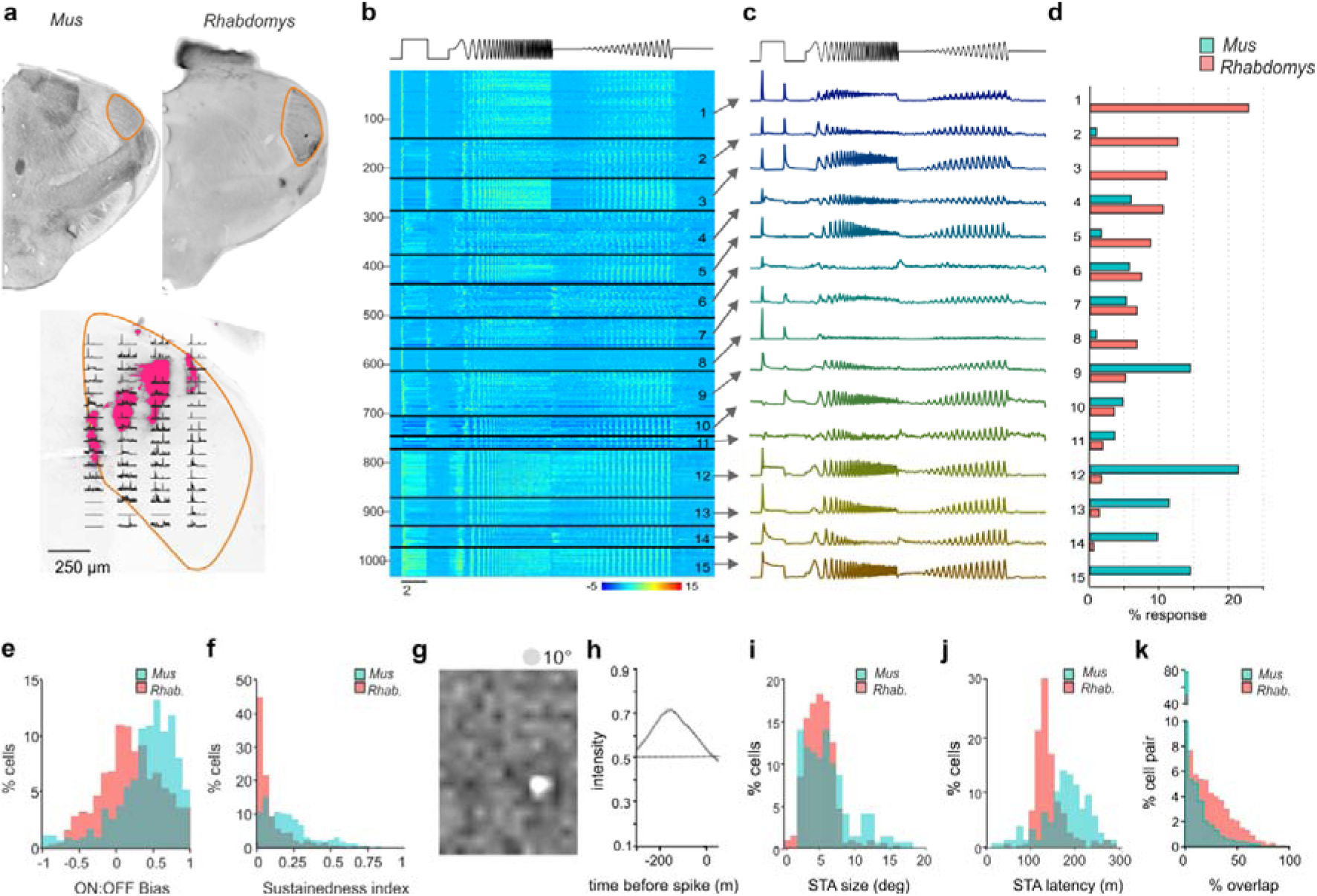
dLGN response diversity in *Mus* and *Rhabdomys*. **a)** Above: coronal sections from *Mus* and *Rhabdomys* with dLGN highlighted with orange edging. Below: Histological reconstruction of recording sites in *Rhabdomys* dLGN. Fluorescence shows diI markings remaining after electrode insertion. Peristimulus time histograms (PSTHs) representing multiunit light-evoked activity recorded at each recording site overlayed (2s light step with 1s pre and post stimulus). All data are scaled according to their maximum firing rate. **b**) Heat map showing normalised PSTHs of dLGN neurons as a function of response cluster (labelled to right), organised as a function of number of *Rhabdomys* responses. Stimulus profile denoted above. **c**) Mean response profile of normalised responses from each cluster (combined across species). Stimulus profile denoted above. **d**) Bar graphs showing proportion of responses within each cluster, for each species (*Mus*: cyan, *Rhabdomys*: red). **e**) Histogram of ON:OFF bias in *Mus* (cyan) and *Rhabdomys* (red). **f**) Histogram of sustainedness index in *Mus* (cyan) and *Rhabdomys* (red). **g&h**) Example spatial **(g**) and temporal (**h**) spike triggered averages recorded in response to binary noise stimulus. **i**) Histogram of receptive field sizes in *Mus* (cyan) and *Rhabdomys* (red); Kolmogorov-Smirnov test P=0.036. **j**) Histogram of receptive field latency (time of peak absolute response) in *Mus* (cyan) and *Rhabdomys* (red); Kolmogorov-Smirnov test: P<0.001. **k**) Histogram of receptive field overlap between pairs of neurons recorded in *Mus* (cyan) and *Rhabdomys* (red); Kolmogorov-Smirnov test: P<0.001.

To explore the characteristics and numbers of response types within each species, we pooled light responsive units from *Rhabdomys* and *Mus,* and extracted isolated response features from the peristimulus histograms (PSTH) of light-responsive neurons using a sparse principal component analysis (sPCA; Supplementary Figure 2; see Methods). We then clustered the data by applying a Gaussian mixture model to the feature set and optimised the cluster number by minimising the Bayesian Information Criteria (Supplementary Figure 2b). This approach produced 15 functional clusters, which varied in their response sign, temporal frequency tuning and/or contrast sensitivity function (Figure 1b-c; Supplementary Figure 3 and 4). Mapping the location of all recorded units across the dLGN ruled out any systematic sampling bias across clusters (Supplementary Figure 1c).

As expected from earlier work, we found response types corresponding to transient ON (clusters 4, 5, 9, 13) and transient OFF (10, 11), sustained ON (12, 14, 15), and ON-OFF responses (1, 2, 3, 7, 8). Within those categories, there was further diversity in temporal and contrast sensitivities. Clusters 1, 3, 5, 9 and 13 showed band-pass temporal frequency tuning, with the remainder showing low-pass tuning. Clusters 5, 13 and 14 showed the highest contrast sensitivity (sensitivity defined by half-maximum of Naka-Rushton curve, see Methods), while 4 and 6 had lowest sensitivity, and clusters 2 and 5 showed saturating contrast sensitivity curves. Suppressed-by-contrast responses were also identified (clusters 6 and 8).

We next compared the proportion of neurons from *Rhabdomys* or *Mus* that were assigned to each functional cluster (Figure 1d). Most clusters (12/15) contained units from both *Mus* and *Rhabdomys*, and where this was the case, the mean response profile remained consistent between species (Supplementary Figure 2H). However, three types appeared to be unique to one species (1, 3 and 15) and frequencies differed by >5-fold for others (2, 5, 9, 12, 13 and 14; Figure 1d). Accordingly, the overall frequency distribution of units across clusters was highly significantly different between species (chi square test, p<0.001). This pattern was consistent across experimental animals and robust to controls for cluster number, batch effects and critical analysis parameters (Supplementary Figure 2C-G).

The 3 most common clusters in *Rhabdomys* (transient ON-OFF; clusters 1-3) were represented by only 4 units in *Mus.* Conversely the clusters with highest *Mus* representation (sustained ON; 12-15) were sparsely represented in *Rhabdomys* (21 units across these 4 clusters). To independently verify these prominent differences between the two species, we separately classified dLGN response units according to the kinetics and polarity of their response to a 2s step (2s 97% contrast step every 20s) (Methods; Figure 1e-f; Supplementary Figure 3). The distribution of units across the resultant categories was again significantly different between species (chi square test p<0.001), confirming that the most common response type was ON-OFF in *Rhabdomys* and sustained ON in *Mus*.

Among the responses to the chirp element of the stimulus >50% of *Rhabdomys* units but only ∼33% of *Mus* units were from clusters displaying band-pass temporal frequency tuning (1, 3, 5, 9, 13). Accordingly, calculations across the whole dLGN population confirmed a preference for higher frequencies in *Rhabdomys* (Supplementary Figure 4). There was a slight but significant shift towards increased contrast sensitivity in *Rhabdomys* compared to *Mus* (Supplementary Figure 4c, d, g and h)*. Mus* neurons were overrepresented in clusters with the lowest contrast sensitivity (5, 13, 14), while *Rhabdomys* neurons predominated clusters with high contrast sensitivity (4, and 6). Clusters showing saturating responses (2 and 5) were also predominant in *Rhabdomys*. Suppressed-by-contrast profiles were detected in both species, albeit at low frequency (clusters 6 and 8).

### *Rhabdomys* dLGN provides higher temporal fidelity and denser coverage of the visual field

To define the spatial information capacity of *Mus* and *Rhabdomys*, we next mapped receptive fields using a binary dense noise stimulus (5Hz). Using spike triggered averages (STAs), we were able to reconstruct receptive fields in 41% and 64% of light-responsive neurons in *Mus* and *Rhabdomys*, respectively. Spatial receptive fields (RFs) had robust centre responses, which were delineated as ON or OFF polarity (Figure 1g-i). Robust opposing surround responses were rare in both species, consistent with previous work in *Mus,* and are not further quantified here. In *Mus,* the mean RF diameter was consistent with prior estimates (median: 5.54;^16,17^; Figure 1i). In *Rhabdomys*, this value was only 6% smaller (median: 5.20; Kolmogorov-Smirnov P=0.036) although the distribution of RF sizes in each species covered a similar range. The *Rhabdomys* dLGN displayed higher temporal fidelity, with the temporal filter in the receptive field centre having significantly shorter latency in this species (median = 134ms and 186ms in *Rhabdomys* and *Mus,* respectively; Kolmogorov-Smirnov test: P<0.001; Figure 1j). Although individual RFs were similar in the two species, the expanded volume of the dLGN allowed greater overlap of RFs in *Rhabdomys* and therefore denser coverage at the population level (Figure 1k).

### Enhanced non-linear spatial integration in *Rhabdomys* dLGN

The higher frequency of ON-OFF units in *Rhabdomys* than *Mus* suggests that visual responses in this species may involve increased non-linear spatiotemporal integration^18^. To assess this directly, we recorded dLGN responses to inverting gratings at 6 spatial frequencies (each at two phases and at 4 orientations; 1Hz; Michelson contrast = 97%) and identified the optimum phase/orientation for each neuron offline (see Methods). Quantifying the change in firing rate as a function of spatial frequency revealed a significant increase in the sensitivity of *Rhabdomys* dLGN neurons to higher spatial frequencies compared to *Mus.* Thus, at their preferred phase/orientation, most *Mus* neurons showed low-pass tuning across the range of spatial frequencies tested, responding up to around 0.6 cpd (Figure 2a,c,d), while most *Rhabdomys* neurons remained responsive to inverting gratings at the highest spatial frequency tested (1.2 cpd; Figure 2b-e), much smaller than the predicted RF size (∼3-8°). Taken together, these results support the preponderance of non-linear spatial summation in *Rhabdomys* neurons, i.e. combining visual signals over space in a non-linear fashion. To further examine this, we quantified the response amplitude to inverting gratings that were larger than the calculated receptive field size of each neuron (identifying the optimal phase and stimulus orientation). We then calculated a linearity index (LI), a ratio of the response at the fundamental frequency of the stimulus (F1, 1Hz), to the second harmonic (F2; 2Hz)) in response to this stimulus in each species (Figure 2f-o). Around 25% of neurons were ‘non-linear’ in *Mus* dLGN (similar to^17^), in contrast with at least 60% in *Rhabdomys*. In both species, non-linear units responded to inverting gratings at spatial frequencies that were smaller than their receptive field sizes (Figure 2 I,j,n,o) whereas linear units did not (Figure 2g, h, l, m).

**Figure 2.**
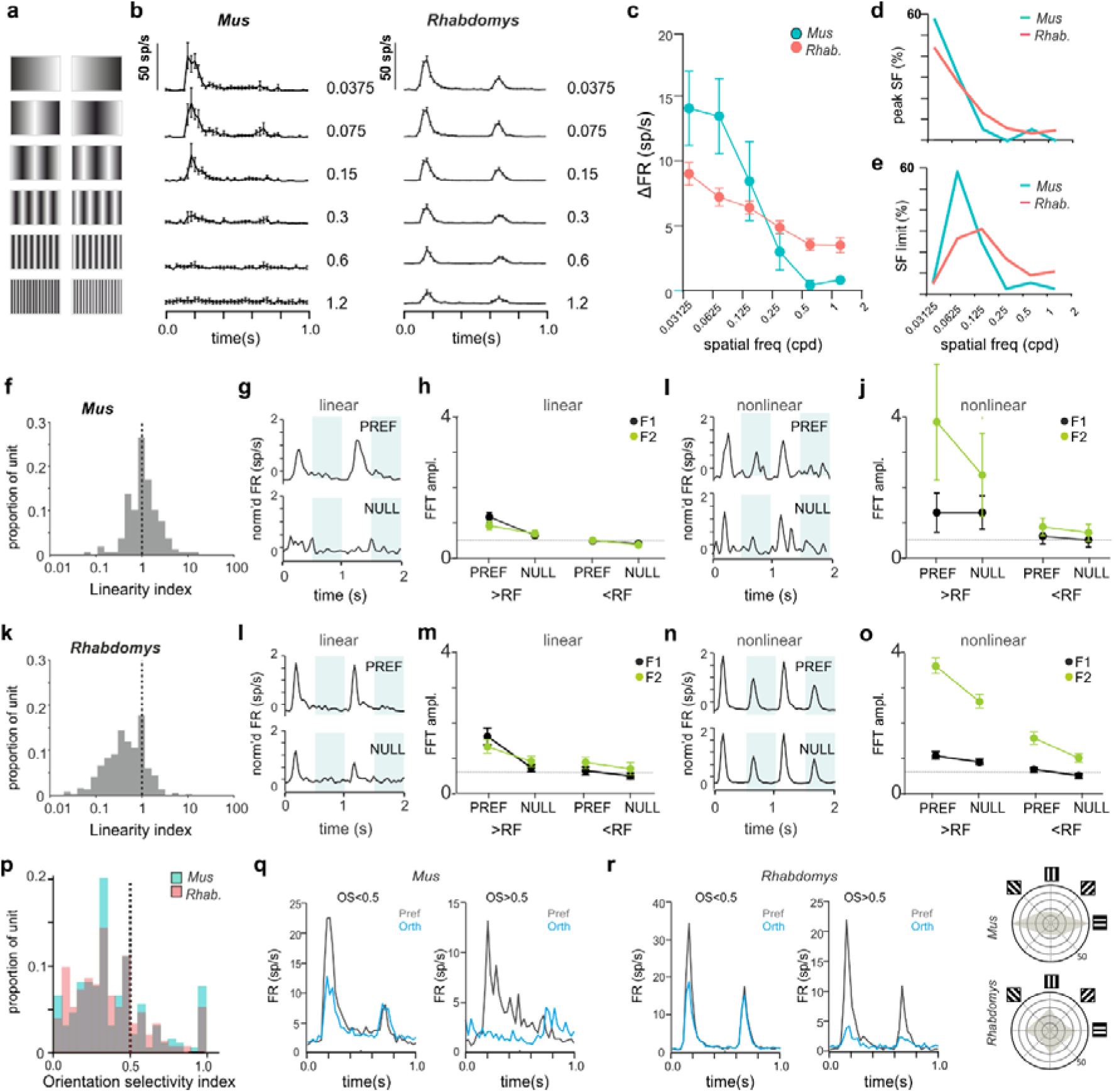
Spatial frequency tuning and linearity in *Mus* and *Rhabdomys*. **a)** Schematic of inverting grating stimuli of increasing spatial frequency. **b**) Mean+/-SEM PSTH for 1Hz inverting grating stimuli at 6 spatial frequencies (0.0375, 0.075, 0.15, 0.3, 0.6, 1.2cpd) for *Mus* (left) and *Rhabdomys* (right) **c**) Amplitude of response to increasing frequency inverting gratings at preferred orientation/phase, shown for *Mus* (cyan) and *Rhabdomys* (red); data show mean+/- SEM. **d&e**) Distribution of % of neurons with peak response at a particular frequency (**d**); and with a threshold to response at a particular spatial frequency (**e**), shown for *Mus* (cyan) and *Rhabdomys* (red). **f,** Distribution of linearity index (LI) in *Mus.* **g,** Mean normalised response of linear neurons in preferred (top panel) or null (lower panel) phase, in response to 2 grating inversions. Shaded regions indicate inversions. **h,** fft amplitude in preferred and null phase of the response, for spatial frequencies greater than (left) or less than (right) the receptive field size, for F1 (black) and F2 (green) frequency components. **i&j,** as in **g&h** but for non-linear responses in *Mus.* **k-o**, as in **f-j**, but for *Rhabdomys.* **p**) Distribution of orientation selectivity index (OSI) for *Mus.* **q&r**) mean response of *Mus* (**q**) and *Rhabdomys* (**r**) neurons with OSI<0.5 (left) and OSI>0.5 (right) in response to inverting grating at optimal spatial frequency, in preferred (black) and null (red) phase. **s,** distribution of preferred orientation of bars for OS neurons (note double plot from 180-360_) for *Mus* and *Rhabdomys* (top and bottom panels, respectively).

Analysis of the inverting grating responses enabled us to determine the degree of orientation selectivity by calculating an orientation selectivity index (OSI; methods) for each neuron at its preferred spatial frequency and phase. The range of OSI values in the *Mus* dLGN was consistent with previous reports^16,17^, and a similar distribution was found in *Rhabdomys* (Figure 2p-r). At a conservative threshold of OSI=0.5, 10% and 11% of light-responsive neurons in *Mus* and *Rhabdomys*, respectively, were classed as orientation selective. In OSI neurons, we found a strong preference for horizontally orientated bars in *Mus,* whereas *Rhabdomys* showed a more even distribution around the tested orientations (Figure 2s); however, the low numbers of OS neurons make it hard to attribute significance to those differences.

### Direction selectivity and responses to motion

We applied a moving bar stimulus to explore motion-sensitivity and direction-selectivity. ∼90% of light-responsive neurons in each species responded to this stimulus with a phasic change in firing. Responsive neurons showed changes in firing at the leading and/or trailing edge of the bar when moving over its receptive field (see representative neurons in Figure 3a,b). Consistent with the evidence of enhanced non-linear spatiotemporal summation in *Rhabdomys*, there was a bias towards responses to both the leading and trailing edge in this species. This could be seen most clearly in the spike-triggered average within an individual neuron’s receptive field (Figure 3c,d). We calculated a direction selectivity index (DSI) for neurons that responded to the moving bar. In a subset of neurons, the strength of responses was dependent on the direction of movement (Figure 3b), consistent with previous reports in *Mus* dLGN^16,17^, however there was a similar distribution of DSIs in both species (Figure 3e). We did not find any statistical difference in the preferred direction of motion between species (Figure 3f; chi square test p<0.001), though *Rhabdomys* showed a bias towards motion sensitivity in the dorso-temporal and ventro-nasal directions.

**Figure 3.**
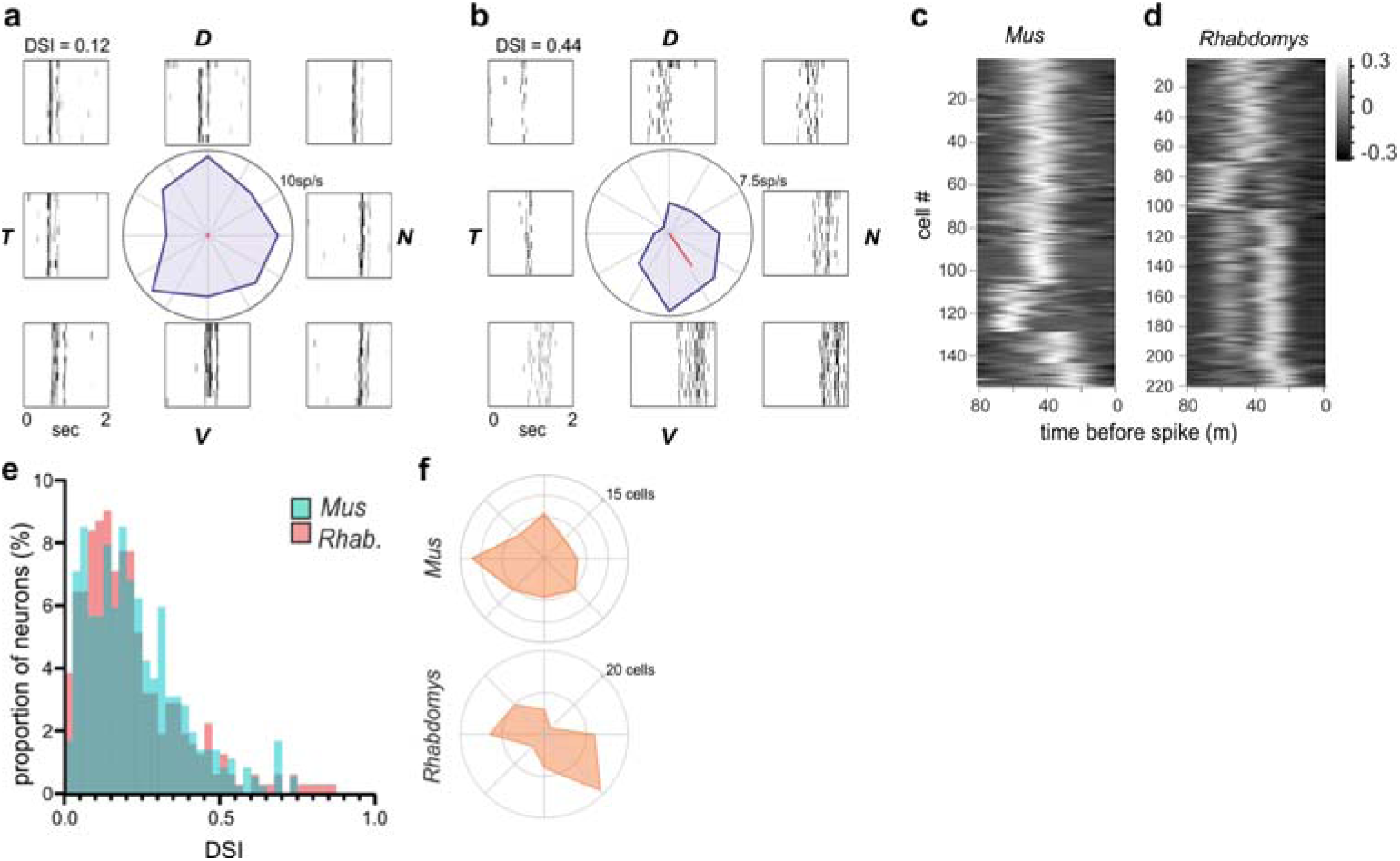
Direction selectivity and responses to motion. **a&b)** Representative neurons from *Rhabdomys* with low (DSI = 0.12) or high (DSI = 0.44) DSI values. Polar plot in centre plots change in firing rate in each direction of motion, and rasters plots to side showing the response to repeated stimuli at each direction. Dorsal, ventral, nasal and temporal locations indicated by D, V, N and T. **c&d)** Heat maps show spike triggered average of luminance changes in the RF centre, as a function of time (−80ms – 0), for all neurons in which we were able to map receptive fields, in *Mus* (c) and *Rhabdomys* (d). **e)** histogram of DSI values for *Rhabdomys* (red) and *Mus* (cyan) dLGN neurons (Kolmogorov-Smirnov test: P=0.193) **f),** distribution of tuning preference for DS neurons in *Mus* (top) and *Rhabdomys* (bottom) with DSI > 0.33.

### Electroretinogram recordings in *Mus* and *Rhabdomys*

While the visual code is sculpted in the dLGN^19^, its fundamental properties are inherited from the retina. Given the marked increase in appearance of OFF excitation in *Rhabdomys* dLGN we therefore turned to *in vivo* electroretinography to determine whether this was also apparent in the retinal light response. Three components of the electroretinogram (ERG) are relevant: the a-wave is derived from photoreceptors, the b wave from ON bipolars and the d wave from OFF bipolars. In response to brief flashes of light under dark-adapted conditions, the a-wave was larger in *Rhabdomys* at high intensities and higher in *Mus* at low intensities, consistent with the relative paucity of rods, which are more sensitive than cones, in *Rhabdomys* (Figure 4A-C). Similarly, the b-wave implicit time (latency to peak), which reflects ON bipolar cell activity, was reduced in *Rhabdomys*, which would be consistent with a reduced contribution of the more sluggish rod pathway to ON responses. Turning to the question of OFF excitation, we recorded ERGs in response to an extended (250ms) step under light adapted (cone isolating) conditions. This stimulus can reveal the activity of ON and OFF cone BCs as separate b- and d-waves associated with the appearance and disappearance of the light step respectively. In accordance with the literature, we found that the light pulse ERG is dominated by the b-wave in *Mus* (Figure 4E-G). Conversely, in *Rhabdomys*, d-waves were at least as prominent (Figure 4E-G) consistent with a strong contribution of OFF bipolar cells and a realignment towards OFF excitation from the earliest step of visual signal transduction in this species.

**Figure 4.**
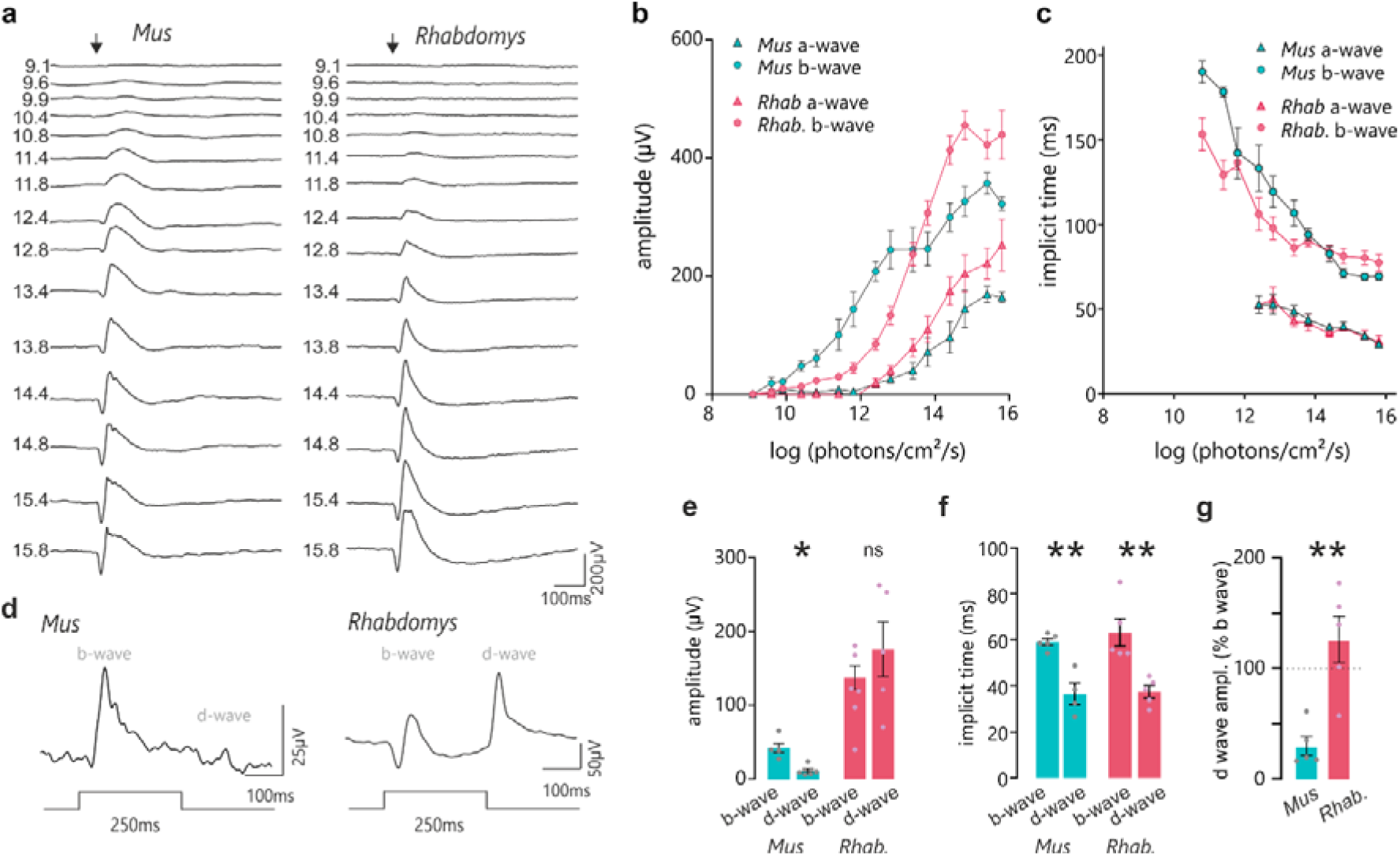
Retinal sensitivity to light in *Mus* and *Rhabdomys*. **a**) Representative traces of dark adapted flash electroretinograms in *Mus* and *Rhabdomys* (left and right, respectively). Flash intensities range from 9.1 to 15.8 log photons/cm^2^/s (shown to left of ERG traces). Arrow indicates flash onset. **b&c)** Amplitude (**b**) and implicit time (**c**) of a and b-waves recorded in *Mus* (cyan) and *Rhabdomys* (red). Data shown mean ± SEM, n=5. **d)** Representative ERG traces recorded from *Mus* and *Rhabdomys* in response to 250ms light step (timecourse indicated below; light intensity 14e-log photons/cm^2^/s). **e**) Quantification of b-wave (light onset) and d-wave amplitudes (light offset) in response to 250ms light step. **f**) Quantification of b-wave and d-wave implicit time in response to 250ms light step. **g**) Quantification of b-wave and d-wave implicit times in response to 250ms light step. d-wave amplitude expressed as % of b-wave amplitude in *Mus* (cyan) and *Rhabdomys* (red). Data shown mean ± SEM, n=5.

### Comparison of *Mus* and *Rhabdomys* retinal cell classes and types

To understand the distinct coding properties in the primary visual pathway of *Rhabdomys* and *Mus* we compared retinal neuronal types of the two species, using atlases derived from single-cell and single-nucleus RNA-seq^20–23^. The *Rhabdomys* atlas^24^ (Figure 5) contained 65,930 nuclei from 2 *Rhabdomys*, which could be classified via standard computational procedures into the five retinal neuronal classes (photoreceptors [PR], horizontal cells [HC], bipolar cells [BC], amacrine cells [AC] and retinal ganglion cells [RGC]) as well as multiple glial types (Muller, Astrocyte, and Microglia), endothelial and retinal pigment epithelial cells (Figure 5a). Each neuronal classes could be further divided into transcriptomically distinct clusters: 3 PR, 1 HC, 18 BC, 33 AC and 33 RGC clusters (Figure 5b). Altogether, the atlas included over 100 retinal clusters, representing putative cell types. Within each class, nearly all *Rhabdomys* clusters mapped in a specific fashion to *Mus* types (see below). This correspondence allowed us to transfer cell type labels of the better-studied *Mus* to *Rhabdomys*, facilitating comparison between the two species.

**Figure 5.**
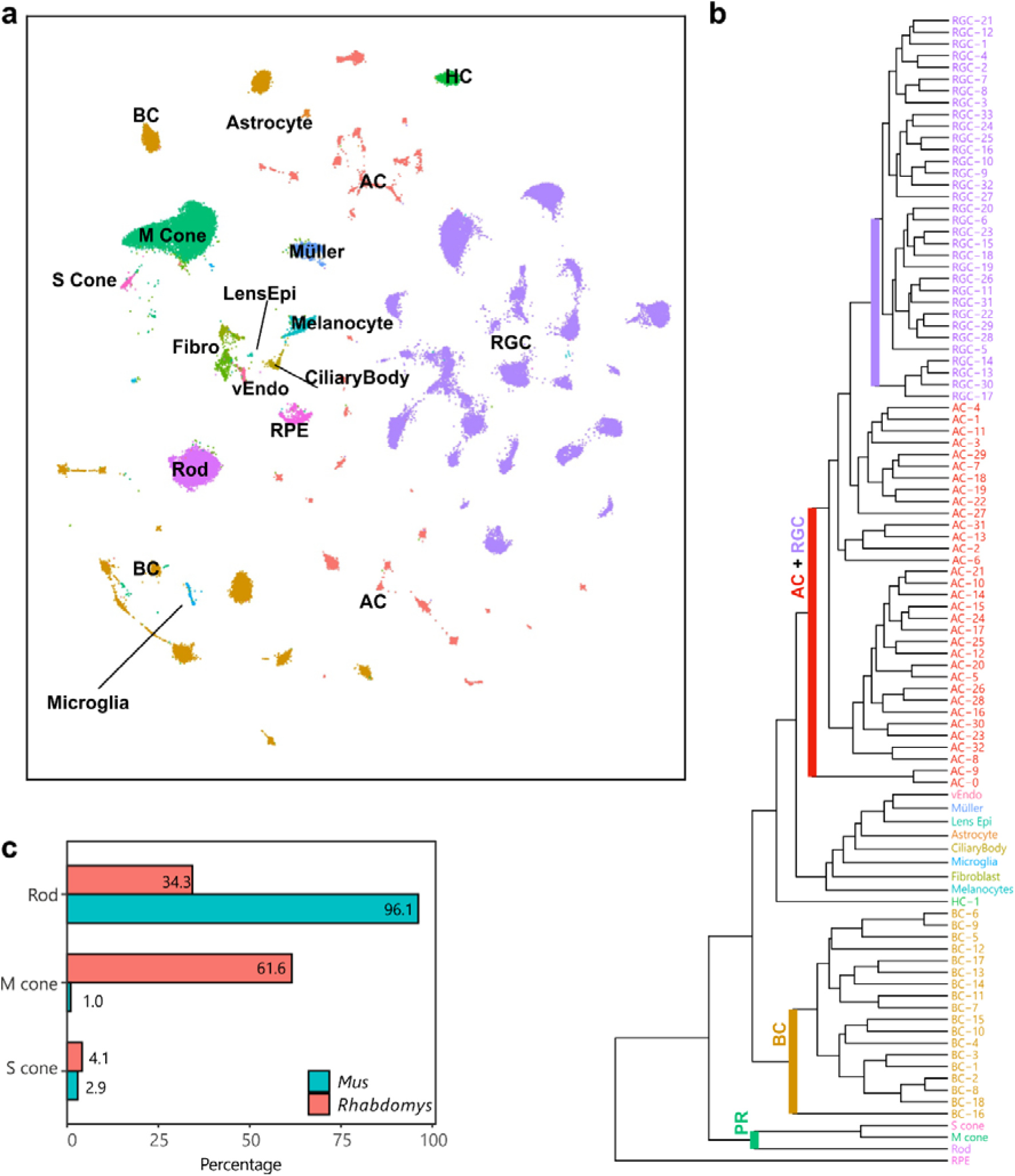
A retinal cell atlas for *Rhabdomys*. **a)** Transcriptionally distinct clusters of *Rhabdomys* retinal cells visualized using UMAP^58^. Cells are coloured by class identity (RPE, retinal pigment epithelial cells; vEndo, vascular endothelial cells; Lens Epi, lens epithelial cells; Fibro, fibroblasts). **b)** Dendrogram showing transcriptional relationships of *Rhabdomys* cell clusters, with major clades corresponding to cell classes. **c)** Bar chart indicating proportion of photoreceptor types in *Rhabdomys* (red) and *Mus* (cyan).

### *Rhabdomys* retina is cone-dominated

Analysis of photoreceptor transcriptomes confirmed and quantified the known shift in rod:cone ratio between *Mus* and *Rhabdomys*: ∼3% of *Mus* photoreceptors but >65% *Rhabdomys* photoreceptors are cones, a >20-fold difference (Figure 5c). Also anticipated was the degree of cone opsin co-expression. ∼40% of *Mus* M cones express S-opsins at low levels^25^, whereas this was true for only ∼1.3% of *Rhabdomys* M cones (112 out of 8622 cones), with the majority (92.7%) expressing only M-cone opsin.

### A reduction in rod BCs and increased proportion of OFF BCs in *Rhabdomys* inner retina

BCs can be subdivided into those that depolarize (ON) or hyperpolarize (OFF) to illumination. Cones innervate both ON and OFF BCs whereas all rod BCs are ON BCs. Clustering the ∼11,400 *Rhabdomys* BC transcriptomes generated 18 putative types (Figure 6a). Based on known markers, these types comprised 7 OFF cone BC types, 9 ON cone BC types, and 1 rod BC type. The final type, BC1B, receives no direct photoreceptor input^20,26^ so cannot be confidently classified as either ON or OFF. A supervised classification analysis based on transcriptomic signatures indicated a predominantly 1:1 correspondence between the *Rhabdomys* BC types and *Mus* BC types (Figure 6b). The patterns of correspondence were consistent with the results of our recent comparative analysis of BC types across 13 mammals using an integrative clustering approach^24^.

**Figure 6.**
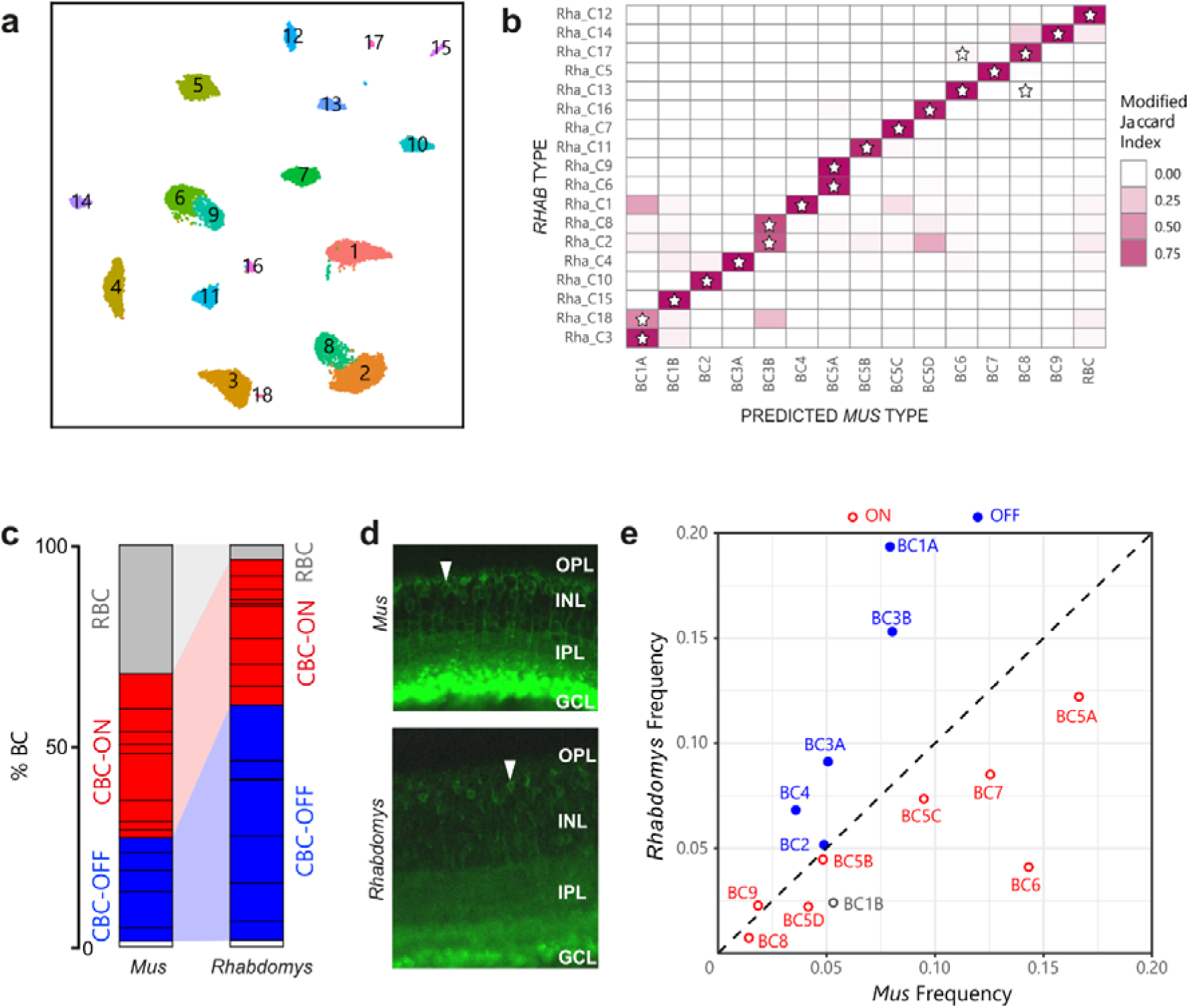
Proportions of bipolar cell types in *Rhabdomys* and *Mus*. **a)** *Rhabdomys* BCs clustered separately and displayed using UMAP. Cells coloured by cluster identity **b)** Confusion matrix indicating the transcriptional correspondence between *Rhabdomys* BC cluster identity (rows) and classifier-assigned *Mus* BC type identity (columns). Cells are coloured based on the modified Jaccard Index (colourbar, right), which ranges from 0 (no-correspondence) to 1 (perfect correspondence; see Methods for details). Pairs of *Rhabdomys* clusters and *Mus* types that belong to the same BC “orthotype” in^24^ are indicated by a star. The preponderance of stars along the diagonal indicates a high concordance between the correspondence analysis presented here and the orthotype analysis in ^24^ **c**) Bar graph showing percentage of BCs assigned rod BC (grey), cone ON BC (red), cone OFF BC (blue) or BC1B (white) cell types in *Mus* and *Rhabdomys*. Dark horizontal lines within each subclass demarcate distinct types. Percentages for biological replicates are shown in Supplementary Table 1. **d**) Transverse sections of *Mus* and *Rhabdomys* stained with antibodies to PKCalpha. INL, Inner nuclear layer; IPL, inner plexiform layer; GCL, ganglion cell layer. RBC axon terminals are labelled intensely; arrowhead indicates an RBC cell body. **e**) Scatter plot comparing the relative frequency of ON (red) and OFF (blue) cone BC types (labelled according to *Mus*) between the two species. BC1B, which cannot be classified as either ON or OFF, is labelled grey.

Three notable differences in BC composition between species were evident. First, the 18 BC types in *Rhabdomys* exceeded the 15 that have been identified and validated in *Mus*. *Mus* types BC1A, 3B and 5A each mapped to two *Rhabdomys* types (Figure 6b). Members of each pair (C3 and C18 for BC1B, C2 and C8 for BC3B, C6 and C9 for BC5A) differentially expressed multiple genes (Supplementary Figure 5), supporting their identity as distinct types. Second, rod BCs comprised around 3.5% of the total BCs in *Rhabdomys*, in contrast to ∼43% in *Mus,* correlating with the difference in rod:cone ratio between species (Figure 6c; Supplementary Table 1). Immunostaining with the RBC marker PKCα validated this difference (Figure 6d). Third, among cone BCs (that is, separate from the difference in rod BC frequency), there are more OFF BCs and fewer ON BCs in *Rhabdomys* than in *Mus*. The four most abundant cone BC types in *Rhabdomys* retina (C1-4; clusters are numbered in order of their abundance) all corresponded to OFF types in *Mus.* Conversely, all but one of the ON cone BC types were more abundant in *Mus* (Figure 6e; Supplementary Table 2). This redistribution of CBC types would predict an enhanced OFF type response at the first retinal synapse, consistent with the enhanced photopic d-wave observed in the ERG (Figure 4).

### Increased abundance of OFF and ON-OFF RGCs in *Rhabdomys*

Reclustering the 26,500 *Rhabdomys* RGCs transcriptomes yielded 33 clusters (Figure 7a), which is substantially lower than the 45 molecularly distinct RGC types in *Mus*. However, supervised classifiers trained on either *Mus* or *Rhabdomys* data indicated that all known 45 *Mus* RGC types were represented among the *Rhabdomys* clusters (Figure 7b). Thus, in several instances a *Rhabdomys* cluster was composed of a group of closely related *Mus* types. The difference in resolution was likely due differences in cell number (26539 in *Rhabdomys* vs 35699 in *Mus*), sequencing depth and/or modality (single-cell vs. single-nucleus).

**Figure 7.**
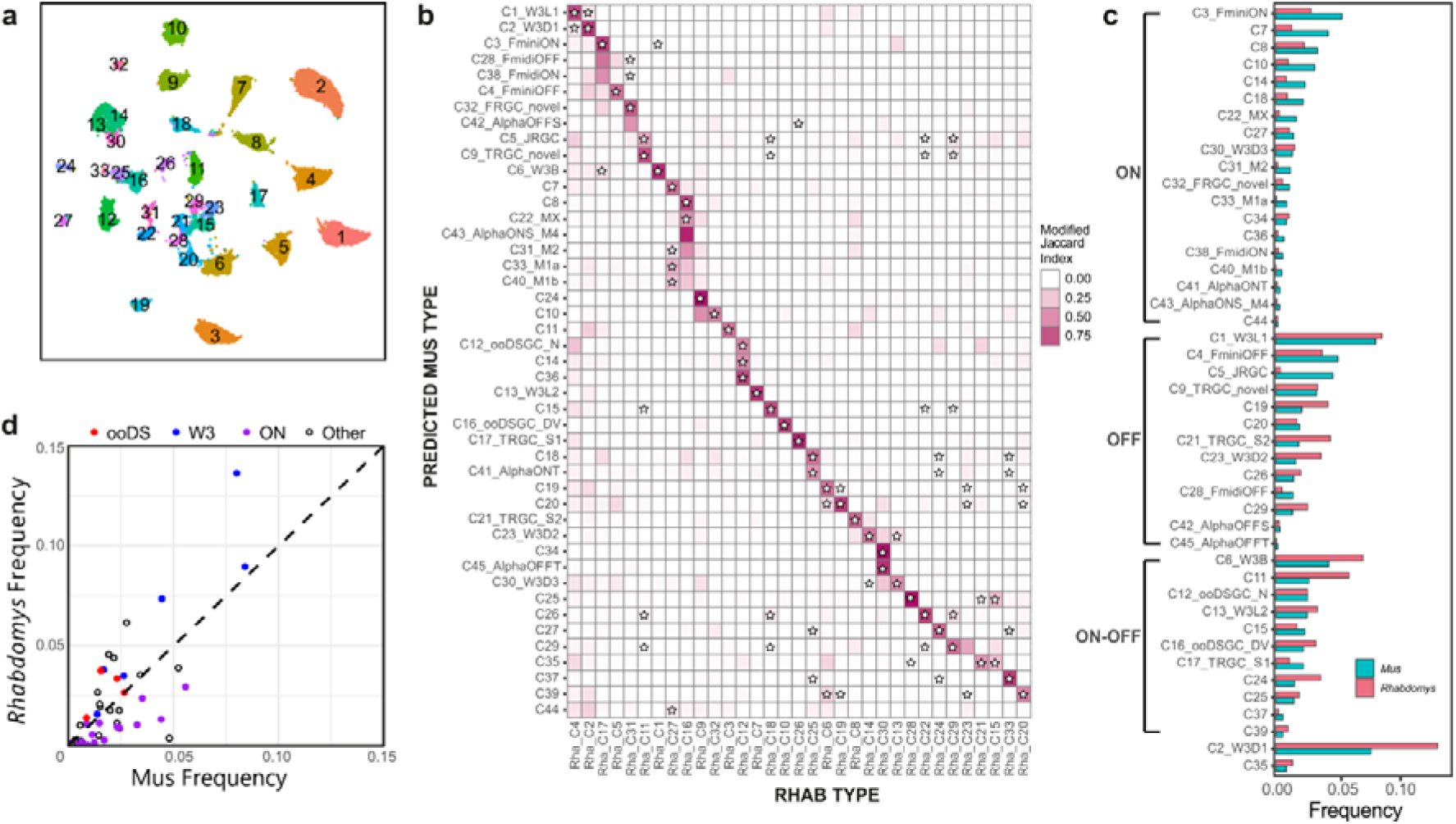
Proportions of RGC types in *Rhabdomys* and *Mus*. **a)** *Rhabdomys* RGCs were clustered separately and displayed using UMAP. Cells are coloured by cluster identity. **b**) Confusion matrix indicating the transcriptional correspondence between *Rhabdomys* RGC cluster identity (columns) and *Mus* RGC type identity (rows). Cells are coloured based on the modified Jaccard Index (colourbar, right), as in panel 6b. Stars indicate pairs of *Rhabdomys* clusters and *Mus* types that belong to the same orthotype in ^24^. This figure shows results of a classifier trained on *Rhabdomys* data. Correspondence was similar when the classifier was trained on *Mus* data. **c**) Bar graph showing the relative frequency of each *Mus* RGC type in both the species. Types are grouped by response polarity – ON, OFF or ON-OFF – based on results in ^21,27–29^. Percentages for biological replicates are shown in Supplementary Table 3. **d)** Scatter plot comparing relative frequency of RGC types between *Rhabdomys* and *Mus*. Response polarity is as shown in **c**. All 6 types of the W3 subclass^21,31^ are more abundant in *Rhabdomys* than *Mus*.

RGCs are conventionally classified as ON, OFF or ON-OFF types depending on whether they are excited by increases or decreases in light intensity or both. Previous studies have combined morphological, physiological, and transcriptomic data to characterize *Mus* RGCs^21,27–29^. Based on those results, we were able to assign 43 *Mus* types to one of these three categories (Figure 7c). Lacking physiological data from *Rhabdomys* RGCs, we used our supervised classification model to provisionally categorize *Rhabdomys* RGCs as ON, OFF or ON-OFF based on their assigned *Mus* label. The 8 most abundant putative ON types in *Mus* were all under-represented in *Rhabdomys*, and most OFF or ON-OFF types were more abundant in *Rhabdomys* than in *Mus*, including all four of the known ON-OFF direction-selective types (ooDSGCs; Figure 7c,d; Supplementary Table 3). This bias is consistent with the preponderance of OFF BCs in retina and OFF and ON-OFF responses in dLGN noted above. Interestingly, orthologues of a subclass of *Mus* RGCs defined by expression in the “THWY3” transgenic line^21,30,31^ were enriched in *Rhabdomys* (Figure 7d); 2 of these types are OFF, 2 ON, 1 ON-OFF, and 1 unknown.

### Depletion of ipRGCs in *Rhabdomys*

Several types of melanopsin-(*Opn4*-) expressing ipRGCs have been characterized in *Mus* ^32–35^. Our *Mus* RGC atlas identifies 5 ipRGC types, which we have provisionally called M1a, M1b, Mx, M2 and M4. Two additional *Mus* RGC types, C7 and C8, are closely related to the ipRGCs; both express the subclass-defining transcription factor *Eome*s, and both express *Opn4* at low levels^21^. *Mus* ipRGC types co-mapped to *Rhabdomys* clusters C16 and C27 (Figure 7b). However, for all these types, the relative abundance was substantially lower in *Rhabdomys* than in *Mus* (Figure 8a), which is the likely reason that they did not form separate clusters.

**Figure 8.**
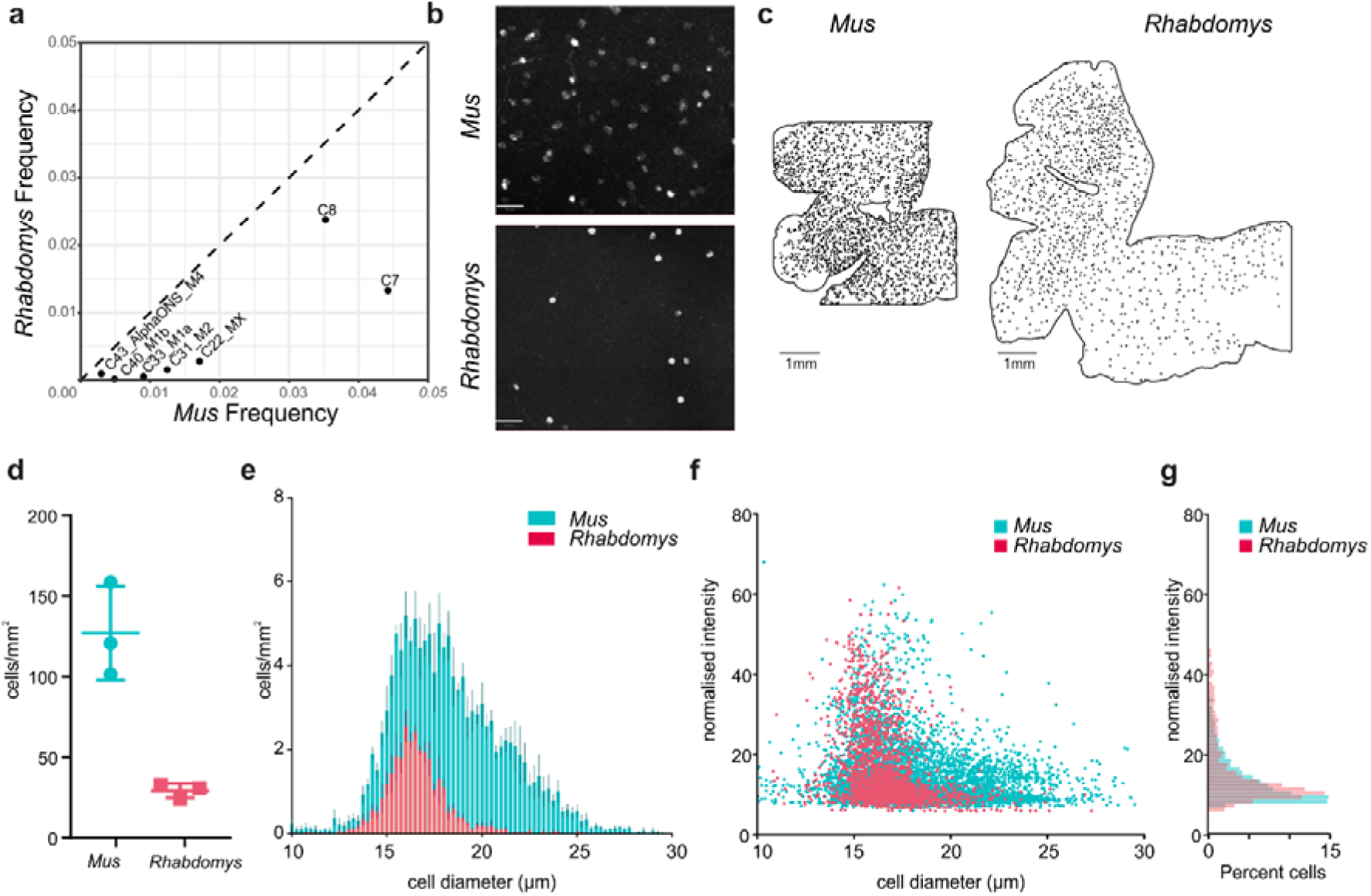
Quantifying ipRGCs in *Rhabdomys* and *Mus* retinae using HCR. **a)** Scatterplot showing the reduced frequency of putative ipRGC types in *Rhabdomys* (y-axis) compared to *Mus* (x-axis). **b**) Representative HCR-FISH of *Mus* and *Rhabdomys* retinal wholemounts, with probe for *Opn4* (species specific). Scale bar = 50µm. **c**) As in b, but showing location of *Opn4* +ve RGCs across whole retina. Scale bar = 1mm. **d**) Number of *Opn4* +ve cells/mm^2^ in *Mus* and *Rhabdomys* retinal wholemounts (n=3; lines show mean±SEM). **e**) Histogram showing diameter of *Opn4* +ve cells/mm^2^ in *Mus* (cyan) and *Rhabdomys* (red) retinal wholemounts (n=3; data show mean±SEM). **f**) Scatter plot showing diameter of *Opn4* +ve cells/mm^2^ vs normalised signal intensity in *Mus* (cyan) and *Rhabdomys* (red) retinal wholemounts (n=3). **g)** Histogram showing normalised intensity of *Opn4* +ve cells in *Mus* (cyan) and *Rhabdomys* (red) retinal wholemounts (n=3 retinae; Kolmogorov-Smirnov test: P<0.001)

To validate these results, we used fluorescent *in situ* hybridisation chain reaction (HCR-FISH) to compare *Opn4+* RGCs in retinal wholemounts from the two species (Figure 8b). Using an automated cell detection method (Supplementary Figure 6), we found that the number of *Opn4*+ cells per unit area was substantially lower in *Rhabdomys* than *Mus* (Figure 8c,d) indicating that both the absolute density of ipRGCs (5x lower) and their representation as a function of the total RGC population were greatly reduced in *Rhabdomys* compared to *Mus*. Quantification of soma size and FISH intensity in the two species (Figure 8e-g) revealed a bias towards cells with smaller soma sizes in *Rhabdomys,* and an increased proportion of cells with high *Opn4* expression.

## Discussion

We applied physiological and transcriptomic techniques to understand how the visual systems of closely related murid rodents are adapted to divergent visual ecology. Diurnal *Rhabdomys* have thicker inner retinas and larger dLGNs than nocturnal *Mus*. Our data reveal that although they apply this extra information capacity to provide denser coverage of the visual scene, this does not amount to simply ‘more of the same’ visual code seen in *Mus*. Rather, we find a marked realignment towards nonlinear spatiotemporal summation in the *Rhabdomys* dLGN. This is apparent in a shift from predominantly ON responses in *Mus* towards biphasic, ON-OFF, responses and increased abundance of units able to track gratings sufficiently fine to fall entirely within their receptive field centre in *Rhabdomys*. These inter-species differences in visual code are accompanied by selective expansions in the abundance of OFF BCs and RGCs with transcriptional fingerprints of ON-OFF response types in *Rhabdomys* compared to *Mus*.

In principle, the enhanced information capacity of the *Rhabdomys* early visual system could have been employed in numerous ways. One option would have been to transmit a higher fidelity, less filtered representation of the scene to the cortex. That would facilitate richer computations and interpretation at higher levels and is the strategy often employed by other species with enhanced visual systems such as primates, where a relatively unprocessed visual code is coupled with an expansion of the visual cortex^36–38^. However, we find that such simple ‘pixel encoders’ (characterised by sustained responses and linear spatial summation) are substantially less common in *Rhabdomys* than *Mus* (Supplementary Figure 3c). At the other extreme, *Rhabdomys*’ extra capacity could plausibly be applied to allow more diverse computations in the early visual system, parsing the visual scene in more complex ways ^39,40^. Our cell atlas analysis provides some support for that proposition, as there is evidence of increased diversity in the BC population with *Mus* subtypes BC1A, 3B and 5A each represented by two transcriptionally distinct clusters in *Rhabdomys*. However, we did not observe a corresponding increase in diversity in the *Rhabdomys* visual code. Thus, unbiased clustering returned around 14 functionally distinct channels in the dLGN of each species (with the proviso that more complexity is expected with a wider array of visual stimuli and/or in distinct RGC targets).

Given the alternatives, what advantage might *Rhabdomys* attain by selectively increasing representation of units with non-linear spatiotemporal summation, especially as such non-linear feature detectors are often more abundant in the retinas of animals that invest less neural resources in (form) vision (e.g.^30^). One possibility is that it represents an efficient approach to enhance spatiotemporal resolution. We find that RF sizes are only marginally finer in *Rhabdomys* compared to *Mus*. With this constraint in mind, natural selection can take two routes to exploit the additional information of *Rhabdomys* to increase spatiotemporal acuity. On the one hand, it can increase the degree of overlap in RF between units to achieve denser coverage of the scene at neuronal population level. We find that this is indeed a feature of the *Rhabdomys* visual code. Alternatively, it can employ non-linear spatiotemporal summation to allow individual neurons to respond to patterns within their RF, achieving greater acuity at the single unit level. *Rhabdomys* appear to exploit this opportunity too, as we find that realignment of the visual code towards non-linear summation allows many units to respond to high frequency gratings.

The primary features of the thalamic visual code are inherited from the retina, and species differences in the visual code that we observe are accompanied by some substantial differences in retinal cellular composition. Most obviously, the *Rhabdomys* retina has many more cone photoreceptors and this is achieved by selective expansion of the M-cone opsin expressing cones (the fraction of photoreceptors expressing S-cone opsin hardly differs between species). The shift towards cones is expected for a more day active species. That this is restricted to M-cone opsin expressing cones is perhaps associated with *Rhabdomys*’ combination of UV sensitive S-cone opsin with a lens that filters out UV-wavelength light^11^.

Second order neurons are also substantially different in *Rhabdomys* with no overlap in the 3 most abundant BC types in each species. The switch to a cone-rich retina in *Rhabdomys* is accompanied by a large decrease in the relative proportion of rod BCs. This may be predicted given the cone bias of the *Rhabdomys* retina, but recent analysis reveals that rod BCs can be numerous even in species with high cone density^24^. Evidence that cones can signal via rod BCs supports the contention that rod BCs do not necessarily become redundant when rod numbers fall. Nevertheless, both anatomical and physiological characteristics of cone BCs facilitate information transfer at higher spatiotemporal resolution.

The decrease in rod BC frequency in *Rhabdomys* compared to *Mus* is accompanied by an increase in cone BC frequency. However, the increase is not distributed equally across cone BC types. Instead, it appears disproportionately in OFF BCs (in order of fractional abundance BC3B, BC4, BC1A, BC3), while cone ON BCs occupy equivalent fractions of the BC population in the two species. The enrichment of OFF BCs in *Rhabdomys* is consistent with the enhanced ERG d-wave amplitude (measuring OFF responses) and the preponderance of OFF excitation responses in the dLGN of this species. A plausible function of the increase in OFF BC frequency facilitates non-linear spatiotemporal summation and, indeed, the two most expanded BC classes in *Rhabdomys* (BC3B and BC4) have characteristics well suited to this purposes (small dendritic and axonal fields and transient responses^41,42^). But realignment towards OFF BCs may bring additional benefits. OFF BCs respond faster than ON BCs, allowing the possibility not only for faster visual reflexes but also higher acuity^43^. As a day active species, *Rhabdomys* may place a higher premium on detecting shadows (attributed to OFF pathways^44^) and in a more general sense the negative spatial contrast to which natural scenes are biased^45^ than high sensitivity vision (attributed to ON pathways^46^). Finally, the enhanced capacity of OFF pathways could support better ability to detect overhead (dark) looming stimuli indicative of aerial predators^47^.

Species differences in the RGC population are more nuanced than those in photoreceptor or BC makeup but are in accordance with the changes in visual code that we observe. Thus, in the *Rhabdomys* atlas RGC types characterised by ON responses in mice are relatively under-represented in favour of those with OFF and ON-OFF responses. The population of ON type, melanopsin-expressing, ipRGCs provides a noteworthy example of this realignment. Not only is the density of *Opn4-*positive RGCs in the *Rhabdomys* retina ∼5-fold less than *Mus*, but their anatomical features suggest a particular reduction in those ipRGCs contributing to thalamocortical vision. Compared to Mus, *Rhabdomys* Opn4+ cells are biased towards smaller soma and slightly higher melanopsin expression. These are characteristics of the M1-3 ipRGCs which subserve reflex responses of circadian entrainment and pupil constriction in Mus. It follows that cells with the morphological characteristics of the *Mus* M4 population^34,48^ that contribute to sustained-ON responses in the dLGN^49–51^ are especially rare in *Rhabdomys*.

The *Mus* vs *Rhabdomys* comparison represents a case study in how sensory systems can adapt to different ecology. Despite their similarity in phylogeny, size and diet these species have evolved markedly different visual systems. *Rhabdomys* is adapted to exploit the higher signal:noise of visual signals in the day with its cone-rich retina and expansion in the thickness of the inner retina and dLGN. We find that it applies this additional information capacity in both predictable and more surprising ways. Thus, in addition to simply achieving greater density of coverage of the visual scene, changes in composition of the retinal cell population drive a realignment in favour of OFF responses and non-linear spatiotemporal summation, which increase spatial resolution of the visual code and may provide other benefits. Our data thus show how evolutionarily advantageous changes in computational outcome can be produced by selective expansion/contraction of cell types comprising a neural circuit.

## Methods

### Animals

Animal care was in accordance with the UK Animals, Scientific Procedures Act of 1986, and the study was approved by the University of Manchester ethics committee. Animals were housed on a 12h:12h light:dark cycle at 22°C with food and water available ad libitum. All experiments were performed in adult *Rhabdomys*□*pumilio* or C57BL6J mice (aged 3–8 months).

### In vivo electrophysiology

*In vivo* electrophysiological recordings were performed in 6 *Rhabdomys* and 5 *Mus* (male), using methods described previously^11^. Anaesthesia was induced with 2% isofluorane in oxygen, and maintained with an intraperitoneal injection of urethane (1.6 g kg−1, 30% w/v; Sigma-Aldrich). A topical mydriatic (tropicamide 1%; Bausch and Lomb) and mineral oil (Sigma-Aldrich) were applied to the left eye prior to recording. After placement into a stereotaxic frame, the skull was exposed and a small hole drilled _∼_2.5 mm posterior and _∼_2.5 mm lateral to bregma (*Rhabdomys*); or _∼_2.3 mm posterior and _∼_2.3 mm lateral to bregma (Mus). A 256-channel recording probe (A4×64-Poly2-5mm-23s-250-177-S256, NeuroNexus Technologies, Inc., Ann Arbor, MI, USA) consisting of 4 shanks spaced 200 µm apart, each with 64 recording sites, was lowered a depth of _∼_3–3.5 mm into the *Rhabdomys* brain, or 2.5-3mm into the *Mus* brain, targeting the dLGN in each species. Broadband neural signals were then acquired using a SmartBox recording system (NeuroNexus Technologies, Inc.), sampling at 20 kHz. Following recordings, data from each of the four electrode shanks were pre-processed by common median referencing, high-pass filtered at 250 Hz and then passed to an automated template-matching-based algorithm for single unit isolation (Kilosort; ^52^). Isolated units were then extracted as virtual tetrode waveforms for validation in Offline Sorter (V3, Plexon, Dallas, TX, USA). Here, unit isolation was confirmed by reference to MANOVA F statistics, J3 and Davies-Bouldin validity metrics and the presence of a distinct refractory period (greater than 1.5 ms) in the interspike interval distribution. Spike sorted data were further analysed in MATLAB R2018a (The MathWorks).

### Visual stimuli

Responses were recorded to a standardised set of temporally and spatially patterned monochromatic stimuli, displayed using an LCD display (width: 26.8cm height: 47.4cm; Hanns-G HE225DPB; Taipei, Taiwan) angled at ∼45° from vertical and placed at a distance of ∼21cm from the contralateral eye to occupy ∼96° x ∼63° visual angle. The temporal stimulus set consisted of a 2s step from minimum to maximum light intensity (98% contrast), followed by 2s of dark, 2s at half maximum light intensity, an 8s temporal chirp (sinusoidal modulation between dark and maximum intensity at 1-8Hz accelerating at rate of 1Hz/s), 2s at half maximum light intensity, and an 8s contrast chirp (sinusoidal modulation at 2Hz increasing from 3% to 97% contrast), as in^15^. Spatial stimuli comprised of a sparse binary noise stimulus (5Hz, square size = 4.2°); inverting grating stimuli (1Hz) at spatial frequencies of 0.03 to 1.2 cpd, presented at two phases and four orientations (0°, 45°, 90° and 135°); and a single bar (4.2°) moving in one of 8 directions (0°, 45°, 90°, 135°, 180°, 225°, 270° and 315°) in a pseudorandom sequence. Stimulus spectra were designed to approximate the activation of each photoreceptor by natural daylight for each species (14.8 MWS effective photons/cm^2^/s; 12.8/12.0 SWS effective photons/cm^2^/s for *Mus* and *Rhabdomys*, respectively; 14.8 rod effective photons/cm^2^/s and 14.7 melanopsin effective photons/cm^2^/s).

### Analysis of dLGN responses to visual stimuli

#### Full field stimuli

Perievent spike histograms (PSTH) were generated with bin size of 30ms. Light responsive units were identified using confidence limits test based on responses to the initial 2s step of the chirp stimulus: units were classified as significant if spike firing rate during the response window was greater than 2 standard deviations above (excitation) or below (inhibition) mean firing rate during baseline window – equivalent to 95% confidence limit. ON:OFF Bias index and Sustainedness index were calculated using previously described methods (Farrow & Masland, 2011; Lindner et al., 2021).

To analyse temporal chirps, the mean response amplitude (maximum – minimum normalised firing rate) for each temporal frequency was fit with a half-gaussian model (^16^) using least-squares minimisation to identify 5 best-fit parameters (low baseline, high baseline, gaussian spread, peak response and peak frequency).

Equation for Half Gaussians is:

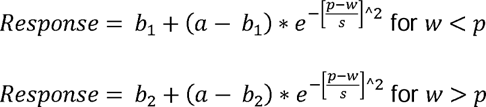

Where *w* is the temporal frequency (Hz), *p* is the temporal frequency (TF) that produces peak response, *a* is the maximum response amplitude at optimum TF, *s* is the Gaussian spread, *b1* is the baseline for frequencies lower than peak TF, *b2* is the baseline for frequencies greater than the peak TF. Peak temporal frequency was rounded to nearest integer to address the limited resolution of temporal frequency analysis (sample every 1Hz).

For contrast chirps, the response amplitude (maximum – minimum normalised firing rate during each period of the contrast chirp stimulus) was normalised to baseline activity (1s before contrast chirp onset) for each unit. This was plotted against Michaelson contrast and then fit using a Naka-Rushton curve using least-squares minimisation to identify 4 best-fit parameters (top, bottom, C50 and slope).

Equation for Naka-Rushton curve is

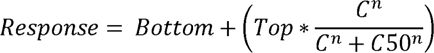

Where *n* = slope, *C* = Michelson contrast and C50 is contrast that produces half maximum response. C50 was constrained between 0 and 1, and slope was constrained between 0 and 10.

#### Functional clustering

Sparse Principal components were generated for the full-field temporal stimulus using the SPaSM toolbox^53^, as described in ^15^. This allows the extraction of response features that are localised in time. We pooled mean PSTH (25ms bins) for all light responsive units from both groups and extracted up to 30 features with 5 non-zero time bins. We then discarded those that accounted for < 1% of the variance. Response features that met these criteria for each window were then combined to produce a total of 30 features for dLGN. sPCs from integrated *Mus* and *Rhabdomys* data were then clustered with a mixture of Gaussian models, a probabilistic model using random initialisation. The optimum number of clusters was determined based on the lowest Bayesian information criteria, which rewards fit but penalises complexity, and a Bayes factor below 6 as a threshold for when there was no longer evidence for further splitting.

To compare distribution of units across communities between two groups, we calculated the distance (Euclidian norm of the difference in the mean relative proportion of neurons in each cluster) between *Mus* and *Rhabdomys* data and compared this with a null distribution obtained by randomly shuffling retinal recordings between two groups 10,000 times. To compare proportion of units from each group within each cluster, we first calculated the % of total units from each group in a given community for each recording, and then found the difference between mean of *Mus* and *Rhabdomys*. This was compared with a null distribution generated as above.

#### Receptive Field mapping

The spatio-temporal receptive field was derived for each unit by generating the spike triggered average (STA) of responses to a sparse binary noise stimulus (5Hz, square size = 4.2°). The separable spatial and temporal components where then extracted from the raw STA matrix by finding the signal peak. RF locations and sizes were then generated by fitting spatial receptive fields with 2D Gaussian function (using lsqcurvefit function, MATLAB). The receptive field size for individual cells was the average of the standard deviation of Gaussians fitted to each dimension. Temporal receptive fields were generated by plotting the temporal response of the RF centre. Receptive field overlap was calculated as a percentage overlap of the 2D Gaussian for pairs of receptive fields recorded in the same animal.

#### Spatial frequency tuning and linearity of spatial summation

To assay changes in spatial frequency tuning, inverting gratings (Michelson contrast between dark and light bars = 98%) were presented in 4 orientations at two phases (phase shifted 90°), at 5 different spatial frequencies (0.03 to 1.2 cpd) at 1Hz. For each unit, response amplitudes were quantified (relative to pre-stimulus firing) for each phase/orientation combination for each spatial frequency to determine the optimal stimulus (R_pref_). R_orth_ was the response to stimuli presented at 90° to the preferred orientation. The orientation selectivity index (OSI) was calculated as the ratio of (R_pref_-R_orth_)/(R_pref_+R_orth_) at the preferred spatial frequency. Cells exceeding an OSI of 0.5 were classed as ‘orientation selective’.

Response linearity was evaluated by quantifying the firing rate of units in response to stimuli that were larger than the calculated receptive field size for an individual unit. Continuous firing rates during the stimulus presentation were Fourier analysed to extract amplitudes of the first and second harmonic components (F1 and F2, at 1Hz and 2Hz), at the preferred and null (90° phase shifted) stimulus (F1_pref_, F1_null_, F2_pref_, F2_null_). A linearity index (LI) of the response was then calculated as F1/F2 for both preferred and null phases, whereby a LI<1 describes a dominant F2 amplitude, indicating nonlinear spatial summation.

#### Motion selectivity

A single drifting bar (4.2°; Michelson contrast= 98%) moving in one of 8 directions (0°, 45°, 90°, 135°, 180°, 225°, 270° and 315°) in a pseudorandom at a speed of. For each unit, response amplitudes were quantified during the presentation of movement (relative to pre-stimulus baseline) for each direction of motion. Since the location of the stimulus relative to the location of a unit’s receptive field was not precisely known, to explore the kinetics of responses, a STA was generated (as described above) for the preferred direction of motion. For presentation purposes, responses were clustered using a Kmeans cluster (MATLAB) finding 3 response clusters.

Two methods were used to explore direction selectivity. First, the mean and variance of circular data were computed using the CircStat, a MATLAB toolbox^54^, to describe the angle and magnitude of directional selectivity. A direction selectivity index was also calculated as the ratio of (Rpref-Rnull)/(Rpref+Rnull), where Rpref was the direction of motion at which the maximum evoked response occurred, and Rnull was response to movement in the opposite direction to this. Cells exceeding a direction selectivity index of 0.33 were classed as ‘direction selective’.

### Electroretinography

ERGs were recorded in adult *Mus* (aged 4-5 months) and adult *Rhabdomys* (aged 7-8 months). All ERGs were recorded at subjective midday following dark adaptation for a period of 6 hours. Animals were anaesthetised under isoflurane in a 95/5% Oxygen/CO2 mix at a flow rate of 0.5 – 1.0L/min. Isoflurane concentrations of 5% and 1.5-3.5% were used for induction and maintenance of anaesthesia, respectively. A topical mydriatic (tropicamide 1%; Bausch and Lomb) and hypromellose eye drops were applied to the recording eye before placement of a corneal contact-lens–type electrode. A needle reference electrode (Ambu, Neuroline) was inserted approximately 5mm from the base of the contralateral eye, and a second subcutaneous needle in the scruff acted as a ground. Electrodes were connected to a Windows PC via a signal conditioner (Model 1902 Mark III, CED) that differentially amplified (X3000) and filtered (band-pass filter cut off 0.5 to 200Hz) the signal, and a digitizer (Model 1401, CED). Core body temperature was maintained at 37°C throughout recording with a homeothermic heat mat (Harvard Apparatus).

Visual stimuli were generated with a combination of violet, blue and cyan elements of a multispectral LED light source (Lumencor). Intensities were modulated via pulse width modulations via an Arduino Uno. Light from the light engine passed through a filter-wheel containing neutral-density filters (reducing the light by between 10^1^ and 10^5^) and focused onto opal diffusing glass (5mm diameter; Edmund Optics Inc.) positioned <5mm from the eye. All LED intensities were controlled dynamically with a PC. Stimuli were measured at the corneal plane using a spectroradiometer (SpectroCAL II, Cambridge Research Systems, UK) between 350-700nm. Dark-adapted stimuli were presented as a 10ms flash of stimulus spectra from background across a range of 9.1 to 15.8 photons/cm^2^/s (interstimulus intervals ranging from 1-6s with increasing intensities). Light-adapted stimuli were presented as square-wave modulations from background at 80.5% Michelson contrast at 2Hz at a background of 14.6 log photons/cm^2^/s, after 15 minutes background adaptation. ERG responses were analysed in MATLAB. For flash responses, a-wave amplitude was calculated relative to baseline prior to stimulus onset, with b-wave with reference to the a-wave trough. For step responses, b- and d-waves were calculated relative to preceding a-wave.

### Immunohistochemistry

Immunostaining of *Mus* and *Rhabdomys* retinal sections and wholemounts was performed as described previously^11^. *Rhabdomys* retinal wholemounts were labelled using rabbit anti-melanopsin antibody (UF006, Advance Targeting Systems, 1:2000) to label ipRGCs. Retinal sections were labelled using rabbit anti-PKCα (ab32376, Abcam, 1:1000) to label rod bipolar cells. Sections and low magnification wholemount retinas were imaged with an Axio Imager D2 upright microscope and captured using a Coolsnap Hq2 camera (Photometrics) through Micromanager software v1.4.23. High magnification images were collected using an inverted LSM 710 laser scanning confocal microscope (Zeiss) and Zen 2009 image acquisition software (Zeiss).

### Fluorescent in situ hybridisation chain reaction

*Mus* (n=3) and *Rhabdomys* (n=3) retinas were dissected and used for HCR^TM^ RNA-FISH (Molecular Instruments) according to the manufacturer protocol for fixed wholemount tissue. Briefly, retinas underwent a series of dehydration and rehydration steps (75%, 50%, 25% methanol solution) and then treated with proteinase K (10 µg/mL). Retinas were pre-hybridized with probe hybridization solution and incubated overnight in probe solution containing custom ordered probes *Mus* or *Rhabdomys Opn4* (2 pmol). Retinas were pre-amplified in amplification buffer and incubated overnight in amplification solution containing hairpins H1 and H2 (30 pmol each; amplification fluorophore 594). Retinas were washed in sodium chloride sodium citrate tween 20 solution and stored at 4 ◦C before imaging. Cells were detected using QuPath^55^ cell detection feature.

### Analysis of Transcriptomic Datasets

#### Alignment and quantification of gene expression

Preprocessing of raw sequencing data was performed using Cellranger (v6.2, 10X Genomics). Sequencing reads were demultiplexed using “cellranger mkfastq” to obtain a separate set of fastq.gz files for each of the 7 samples. These files were then aligned to a reference genome ^13^ using “cellranger count” with the --include-introns flag to include both exonic and intronic reads, resulting in a gene expression matrix (GEM) summarizing transcript counts within each sample. GEMs from each of the 7 samples were combined (column-wise concatenated) to yield a total GEM.

#### Segregation of major retinal cell classes

Analysis of the total GEM was performed in R, with the workflow based on Seurat v4.3.0 for single-cell analysis developed and maintained by the Satija laboratory ^56^ (https://satijalab.org/seurat/). Transcript counts in each cell were normalized to a total library size of 10,000 and log-transformed (X_→_log_(X_+_1)). We identified the top 2,000 highly variable genes and applied principal component analysis (PCA) to obtain a linear factorization of the submatrix corresponding to these highly variable genes. Using the top 20 principal components for each cell, we built a k-nearest neighbor graph on the data, and then clustered with a resolution parameter of 0.5 using Seurat’s FindClusters function. Each cluster of cells was assigned to a retinal cell class based on canonical markers characterized in mice^22^; for example, *Vsx2*, *Otx2* and *Grik1* were used to identify bipolar cells, and *Rbpms*, *Nefl* and *Nefm* were used to identify RGCs.

#### Supervised classification analysis of transcriptional correspondence between Rhabdomys and Mouse types

RGCs were separated from the *Rhabdomys* total GEM and, as the *Rhabdomys* genome was annotated with mammalian orthologs across murid genome assemblies, were merged with a reference *Mus* RGC atlas^21^ using genes that were present in both the *Rhabdomys* GEM and *Mus* reference GEM to create a joint GEM. We used the Canonical Correlation Analysis-based framework in Seurat, integrating by species of origin, to produce an integrated version of the joint GEM adjusted to account for species specific differences in gene expression. The top 2000 variable features of the integrated GEM were used to train a gradient boosted decision tree using the reference *Mus* RGC cells, implemented in R using the xgboost package^57^. The *Mus* RGC-trained classifier was used to assign a *Mus* identity to each *Rhabdomys* RGC based on its expression of the 2000 training features. To identify how rare *Mus* RGC types mapped to *Rhabdomys* types, a similar decision tree model was trained using the *Rhabdomys* RGC cells, and applied to *Mus* RGCs to assign a *Rhabdomys* identity to each *Mus* RGC. Similarly, this was done with *Rhabdomys* bipolar cells with a relevant *Mus* bipolar cell atlas to assign *Mus* bipolar labels to each *Rhabdomys* bipolar cell, and vice versa. These reciprocal mappings helped us validated the robustness of the label transfer procedure.

To summarize the results of the supervised classification mapping, we calculated a modified Jaccard index for each pair of *Rhabdomys* and assigned *Mus* types. For a given *Rhabdomys* type A and assigned *Mus* type B, we calculated

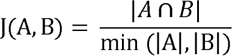

## Data and Code Availability

Analysis scripts for the *Rhabdomys* snRNA-seq data is available via Zenodo (https://zenodo.org/record/8067826) and on our Github (https://github.com/shekharlab/RetinaEvolution). The raw and processed sequencing data produced in this work are available via the Gene Expression Omnibus (GEO) under accession number GSE237210.

## Acknowledgements

This work was supported by a Sir Henry Dale Fellowship 218556/Z/19/Z, jointly funded by the Wellcome Trust and the Royal Society (AEA); a Wellcome Investigator Award 210684/Z/18/Z (RJL); NIH grant U01MH105960 (JRS); and the McKnight Foundation (KS).

## Author contributions

AEA, JS, KS and RJL supervised the project; AEA performed electrophysiological data collection and analysis, with contributions from JM, JR, PO and RS; AEA performed ERG data collection and analysis with contributions from BBO; RR, AP and AEA performed HCR-FISH experiments and analysis; CW and JW performed retinal immunohistochemistry, with contributions from NM; JH performed computational analysis of scRNA-seq data with contribution from WY. AM performed scRNA-seq experiments. AEA, JH, JS, KS and RJL wrote manuscript with input and approval of all authors.

## References

1 Butler, A. B., Hodos, W., ProQuest & ProQuest Ebook, C. Comparative vertebrate neuroanatomy : evolution and adaptation. (2005).

2 Buzsaki, G. et al. Tools for probing local circuits: high-density silicon probes combined with optogenetics. Neuron 86, 92–105, doi:10.1016/j.neuron.2015.01.028 (2015).

3 Jun, J. J. et al. Fully integrated silicon probes for high-density recording of neural activity. Nature 551, 232–236, doi:10.1038/nature24636 (2017).

4 Roberts, R. J. V., Pop, S. & Prieto-Godino, L. L. Evolution of central neural circuits: state of the art and perspectives. Nat Rev Neurosci 23, 725–743, doi:10.1038/s41583-022-00644-y (2022).

5 Tanay, A. & Sebe-Pedros, A. Evolutionary cell type mapping with single-cell genomics. Trends Genet 37, 919–932, doi:10.1016/j.tig.2021.04.008 (2021).

6 Zeng, H. & Sanes, J. R. Neuronal cell-type classification: challenges, opportunities and the path forward. Nat Rev Neurosci 18, 530–546, doi:10.1038/nrn.2017.85 (2017).

7 Chalupa, L. M. et al. Eye, Retina, and Visual System of the Mouse. (2008).

8 Dewsbury, D. A. & Dawson, W. W. African 4-Striped Grass Mice (Rhabdomys-Pumilio), a Diurnal-Crepuscular Muroid Rodent, in the Behavioral Laboratory. Behav Res Meth Instr 11, 329–333, doi:Doi 10.3758/Bf03205671 (1979).

9 Schumann, D. M., Cooper, H. M., Hofmeyr, M. D. & Bennett, N. C. Circadian rhythm of locomotor activity in the four-striped field mouse, Rhabdomys pumilio: A diurnal African rodent. Physiol Behav 85, 231–239, doi:10.1016/j.physbeh.2005.03.024 (2005).

10 van der Merwe, I. et al. The topography of rods, cones and intrinsically photosensitive retinal ganglion cells in the retinas of a nocturnal (Micaelamys namaquensis) and a diurnal (Rhabdomys pumilio) rodent. PLoS One 13, e0202106, doi:10.1371/journal.pone.0202106 (2018).

11 Allen, A. E. et al. Spectral sensitivity of cone vision in the diurnal murid Rhabdomys pumilio. J Exp Biol 223, doi:10.1242/jeb.215368 (2020).

12 Schumann, D. M., Cooper, H. M., Hofmeyr, M. D. & Bennett, N. C. Light-induced Fos expression in the suprachiasmatic nucleus of the four-striped field mouse, Rhabdomys pumilio: A southern African diurnal rodent. Brain Res Bull 70, 270–277, doi:10.1016/j.brainresbull.2006.04.009 (2006).

13 Richardson, R. et al. The genomic basis of temporal niche evolution in a diurnal rodent. Curr Biol 33, 3289–3298 e3286, doi:10.1016/j.cub.2023.06.068 (2023).

14 Morrow, A., Smale, L., Meek, P. D. & Lundrigan, B. Tradeoffs in the sensory brain between diurnal and nocturnal rodents. Brain Behav Evol, doi:10.1159/000538090 (2024).

15 Baden, T. et al. The functional diversity of retinal ganglion cells in the mouse. Nature 529, 345–350, doi:10.1038/nature16468 (2016).

16 Grubb, M. S. & Thompson, I. D. Quantitative characterization of visual response properties in the mouse dorsal lateral geniculate nucleus. J Neurophysiol 90, 3594–3607, doi:10.1152/jn.00699.2003 (2003).

17 Piscopo, D. M., El-Danaf, R. N., Huberman, A. D. & Niell, C. M. Diverse visual features encoded in mouse lateral geniculate nucleus. J Neurosci 33, 4642–4656, doi:10.1523/JNEUROSCI.5187-12.2013 (2013).

18 Enroth-Cugell, C. & Robson, J. G. The contrast sensitivity of retinal ganglion cells of the cat. J Physiol 187, 517–552, doi:10.1113/jphysiol.1966.sp008107 (1966).

19 Roman Roson, M., et al. Mouse dLGN Receives Functional Input from a Diverse Population of Retinal Ganglion Cells with Limited Convergence. Neuron 102, 462–476 e468, doi:10.1016/j.neuron.2019.01.040 (2019).

20 Shekhar, K. et al. Comprehensive Classification of Retinal Bipolar Neurons by Single-Cell Transcriptomics. Cell 166, 1308–1323 e1330, doi:10.1016/j.cell.2016.07.054 (2016).

21 Tran, N. M. et al. Single-Cell Profiles of Retinal Ganglion Cells Differing in Resilience to Injury Reveal Neuroprotective Genes. Neuron 104, 1039–1055 e1012, doi:10.1016/j.neuron.2019.11.006 (2019).

22 Macosko, E. Z. et al. Highly Parallel Genome-wide Expression Profiling of Individual Cells Using Nanoliter Droplets. Cell 161, 1202–1214, doi:10.1016/j.cell.2015.05.002 (2015).

23 Yan, W. et al. Mouse Retinal Cell Atlas: Molecular Identification of over Sixty Amacrine Cell Types. J Neurosci 40, 5177–5195, doi:10.1523/JNEUROSCI.0471-20.2020 (2020).

24 Hahn, J. et al. Evolution of neuronal cell classes and types in the vertebrate retina. bioRxiv, doi:10.1101/2023.04.07.536039 (2023).

25 Applebury, M. L. et al. The murine cone photoreceptor: a single cone type expresses both S and M opsins with retinal spatial patterning. Neuron 27, 513–523, doi:10.1016/s0896-6273(00)00062-3 (2000).

26 Della Santina, L. et al. Glutamatergic Monopolar Interneurons Provide a Novel Pathway of Excitation in the Mouse Retina. Curr Biol 26, 2070–2077, doi:10.1016/j.cub.2016.06.016 (2016).

27 Goetz, J. et al. Unified classification of mouse retinal ganglion cells using function, morphology, and gene expression. Cell Rep 40, 111040, doi:10.1016/j.celrep.2022.111040 (2022).

28 Rousso, D. L. et al. Two Pairs of ON and OFF Retinal Ganglion Cells Are Defined by Intersectional Patterns of Transcription Factor Expression. Cell Rep 15, 1930–1944, doi:10.1016/j.celrep.2016.04.069 (2016).

29 Huang, W. et al. Linking transcriptomes with morphological and functional phenotypes in mammalian retinal ganglion cells. Cell Rep 40, 111322, doi:10.1016/j.celrep.2022.111322 (2022).

30 Zhang, Y., Kim, I. J., Sanes, J. R. & Meister, M. The most numerous ganglion cell type of the mouse retina is a selective feature detector. Proc Natl Acad Sci U S A 109, E2391–2398, doi:10.1073/pnas.1211547109 (2012).

31 Laboulaye, M. A., Duan, X., Qiao, M., Whitney, I. E. & Sanes, J. R. Mapping Transgene Insertion Sites Reveals Complex Interactions Between Mouse Transgenes and Neighboring Endogenous Genes. Front Mol Neurosci 11, 385, doi:10.3389/fnmol.2018.00385 (2018).

32 Hattar, S. et al. Central projections of melanopsin-expressing retinal ganglion cells in the mouse. J Comp Neurol 497, 326–349, doi:10.1002/cne.20970 (2006).

33 Ecker, J. L. et al. Melanopsin-expressing retinal ganglion-cell photoreceptors: cellular diversity and role in pattern vision. Neuron 67, 49–60, doi:10.1016/j.neuron.2010.05.023 (2010).

34 Berson, D. M., Castrucci, A. M. & Provencio, I. Morphology and mosaics of melanopsin-expressing retinal ganglion cell types in mice. J Comp Neurol 518, 2405–2422, doi:10.1002/cne.22381 (2010).

35 Estevez, M. E. et al. Form and function of the M4 cell, an intrinsically photosensitive retinal ganglion cell type contributing to geniculocortical vision. J Neurosci 32, 13608–13620, doi:10.1523/JNEUROSCI.1422-12.2012 (2012).

36 Dacey, D. M., Peterson, B. B., Robinson, F. R. & Gamlin, P. D. Fireworks in the primate retina: in vitro photodynamics reveals diverse LGN-projecting ganglion cell types. Neuron 37, 15–27, doi:10.1016/s0896-6273(02)01143-1 (2003).

37 Van Essen, D. C., Newsome, W. T. & Maunsell, J. H. The visual field representation in striate cortex of the macaque monkey: asymmetries, anisotropies, and individual variability. Vision Res 24, 429–448, doi:10.1016/0042-6989(84)90041-5 (1984).

38 Van Essen, D. C., Anderson, C. H. & Felleman, D. J. Information processing in the primate visual system: an integrated systems perspective. Science 255, 419–423, doi:10.1126/science.1734518 (1992).

39 Jun, N. Y., Field, G. D. & Pearson, J. M. Efficient coding, channel capacity, and the emergence of retinal mosaics. Adv Neural Inf Process Syst 35, 32311–32324 (2022).

40 Ocko, S., Lindsey, J., Ganguli, S. & Deny, S. The emergence of multiple retinal cell types through efficient coding of natural movies. Advances in Neural Information Processing Systems, 9389-9400 (2018).

41 Vielma, A. H. & Schmachtenberg, O. Electrophysiological fingerprints of OFF bipolar cells in rat retina. Sci Rep 6, 30259, doi:10.1038/srep30259 (2016).

42 Euler, T., Haverkamp, S., Schubert, T. & Baden, T. Retinal bipolar cells: elementary building blocks of vision. Nat Rev Neurosci 15, 507–519, doi:10.1038/nrn3783 (2014).

43 Mazade, R., Jin, J., Pons, C. & Alonso, J. M. Functional Specialization of ON and OFF Cortical Pathways for Global-Slow and Local-Fast Vision. Cell Rep 27, 2881–2894 e2885, doi:10.1016/j.celrep.2019.05.007 (2019).

44 Westo, J. et al. Retinal OFF ganglion cells allow detection of quantal shadows at starlight. Curr Biol 32, 2848–2857 e2846, doi:10.1016/j.cub.2022.04.092 (2022).

45 Tadmor, Y. & Tolhurst, D. J. Calculating the contrasts that retinal ganglion cells and LGN neurones encounter in natural scenes. Vision Res 40, 3145–3157, doi:10.1016/s0042-6989(00)00166-8 (2000).

46 Smeds, L. et al. Paradoxical Rules of Spike Train Decoding Revealed at the Sensitivity Limit of Vision. Neuron 104, 576–587 e511, doi:10.1016/j.neuron.2019.08.005 (2019).

47 Yilmaz, M. & Meister, M. Rapid innate defensive responses of mice to looming visual stimuli. Curr Biol 23, 2011–2015, doi:10.1016/j.cub.2013.08.015 (2013).

48 Schmidt, T. M. et al. A role for melanopsin in alpha retinal ganglion cells and contrast detection. Neuron 82, 781–788, doi:10.1016/j.neuron.2014.03.022 (2014).

49 Brown, T. M. et al. Melanopsin contributions to irradiance coding in the thalamo-cortical visual system. PLoS Biol 8, e1000558, doi:10.1371/journal.pbio.1000558 (2010).

50 Allen, A. E., Storchi, R., Martial, F. P., Bedford, R. A. & Lucas, R. J. Melanopsin Contributions to the Representation of Images in the Early Visual System. Curr Biol 27, 1623–1632 e1624, doi:10.1016/j.cub.2017.04.046 (2017).

51 Brown, T. M. et al. Melanopsin-based brightness discrimination in mice and humans. Curr Biol 22, 1134–1141, doi:10.1016/j.cub.2012.04.039 (2012).

52. Pachitariu, M., Steinmetz, N., Kadir, S., Carandini, M. & Harris, K. D. Kilosort:realtime spike-sorting for extracellular electrophysiology with hundreds of channels. BioRxiv (2016).

53 Sjöstrand, K., Clemmensen, L. H., Larsen, R., Einarsson, G. & Ersbøll, B. SpaSM: A MATLAB Toolbox for Sparse Statistical Modeling. Journal of Statistical Software 84, 1–37, doi:10.18637/jss.v084.i10 (2018).

54 Berens, P. CircStat: A MATLAB Toolbox for Circular Statistics. Journal of Statistical Software 31, 1–21, doi:10.18637/jss.v031.i10 (2009).

55 Bankhead, P. et al. QuPath: Open source software for digital pathology image analysis. Sci Rep 7, 16878, doi:10.1038/s41598-017-17204-5 (2017).

56 Stuart, T. et al. Comprehensive Integration of Single-Cell Data. Cell 177, 1888–1902 e1821, doi:10.1016/j.cell.2019.05.031 (2019).

57 Chen, T. & Guestrin, C. XGBoost: A Scalable Tree Boosting System. Proceedings of the 22^nd^ ACM SIGKDD International Conference on Knowledge Discovery and Data Mining 785–794 (2016).

58 McInnes, L., Healy, J. & Melville, J. Umap: Uniform manifold approximation and projection for dimension reduction. arXiv preprint arXiv:1802.03426 (2018).

