## Supplementary Figures and Tables for "Reconfiguration of the visual code and retinal cell type complement in closely related diurnal and nocturnal mice"

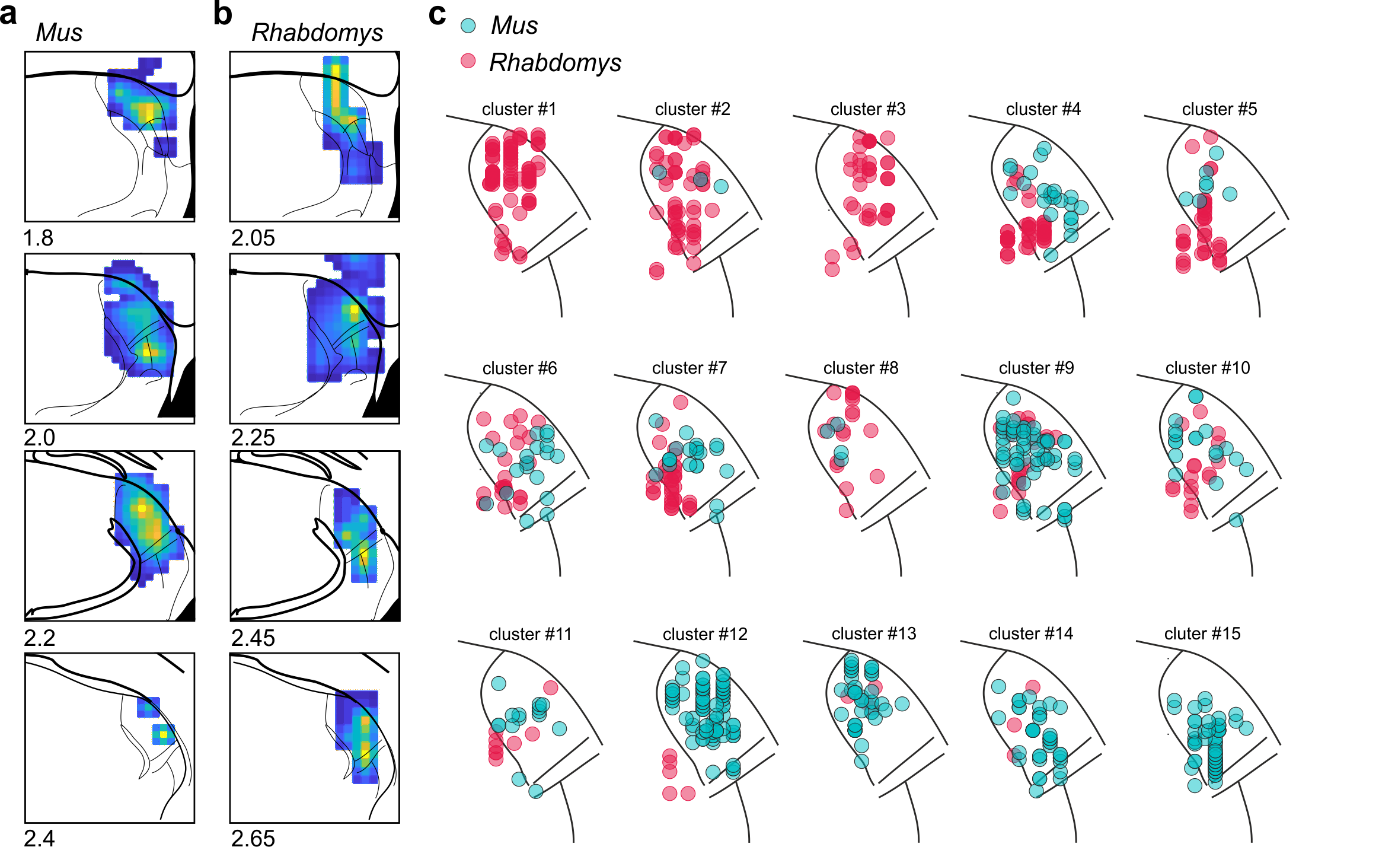

Supplementary Figure 1

**Location of recorded units in *Mus* and *Rhabdomys***

**a&b,** Heat map showing reconstruction of light-evoked responses in *Mus* (a) and *Rhabdomys* (b). Note that histological location of dLGN is approximated in *Rhabdomys* from a scaled version of *Mus* stereotaxic atlas. Numbers below each plot indicate distance from bregma. **c)** Location of units assigned to each functional cluster for *Mus* (cyan) and *Rhabdomys* (red). For simplicity, location is compressed on rostro-caudal axis and superimposed onto dLGN map in coronal plane.

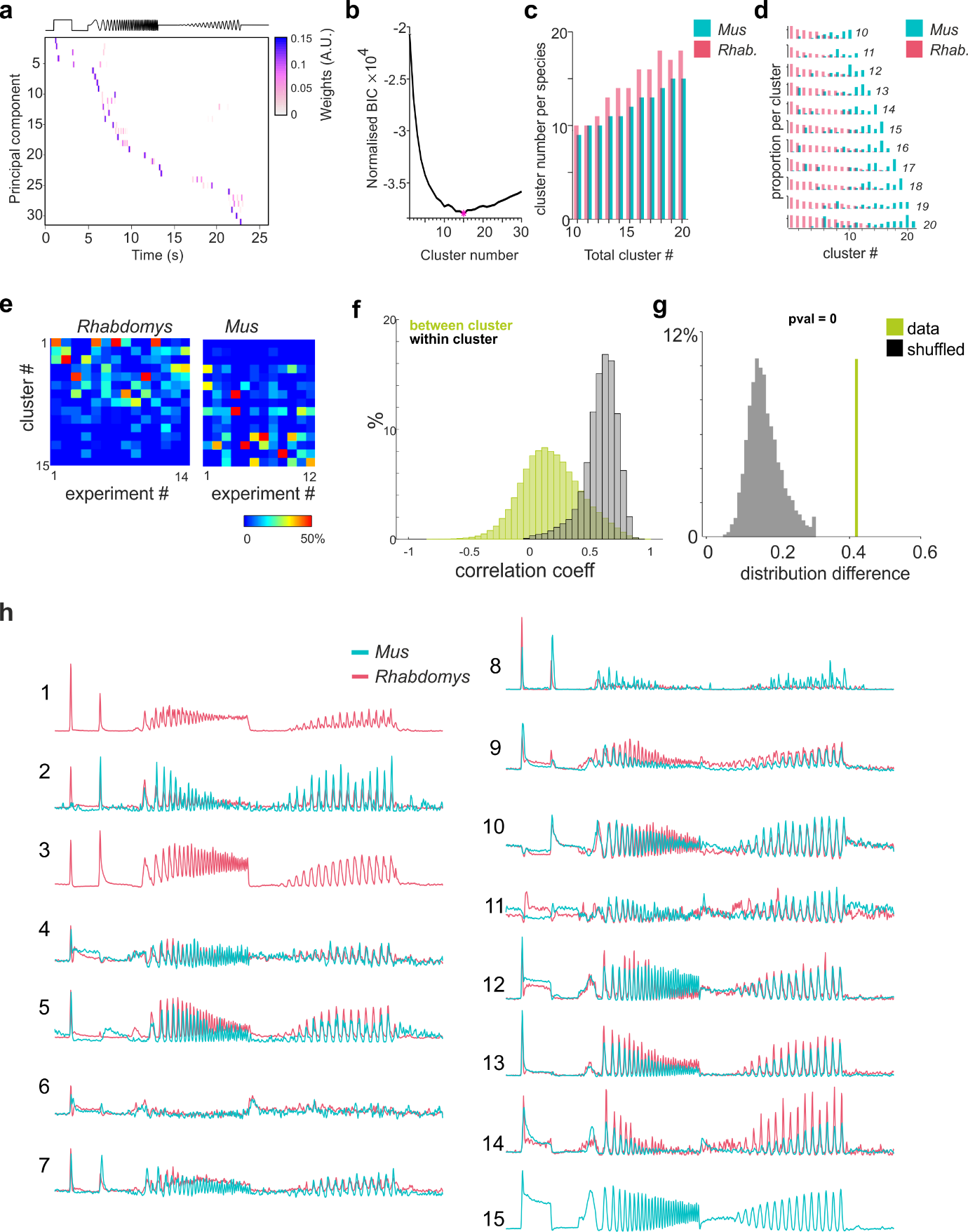

Supplementary Figure 2

**Clustering of visually evoked responses in *Mus* and *Rhabdomys***

**a)** Heatmap showing weights of the principal components from sparse principal component analysis of *Mus* and *Rhabdomys* dLGN unit firing rates in response to temporally modulated stimulus. Stimulus profile is shown in upper panel. Rows of heatmap depict the sparse principal components contributing to >1% variance, organised according to time of peak. **b)** Normalized Bayesian information criterion (BIC) curve; pink point indicates the optimal numbers of clusters (15). **c&d)** Number (c) and distribution (d) of clusters assigned to each species when cluster number is fixed (ranging from 10-20 clusters). Note that irrespective of cluster number, the differences among species remains consistent. *Rhabdomys* in red and *Mus* in cyan. **e)** Heatmap showing percentage of responses from each cluster as a function of experiment (14 recordings in *Rhabdomys*; 12 recordings in Mus). **f)** Histogram showing correlation between *Mus* and *Rhabdomys* cluster PSTHs within clusters (black) and between clusters (red). **g)** Euclidian difference in cluster distribution between species (green line), showed alongside histogram of null distribution of shuffled experiments (p=0). **h**) Mean response profile of each cluster, separated into *Rhabdomys* (red) and *Mus* (cyan).

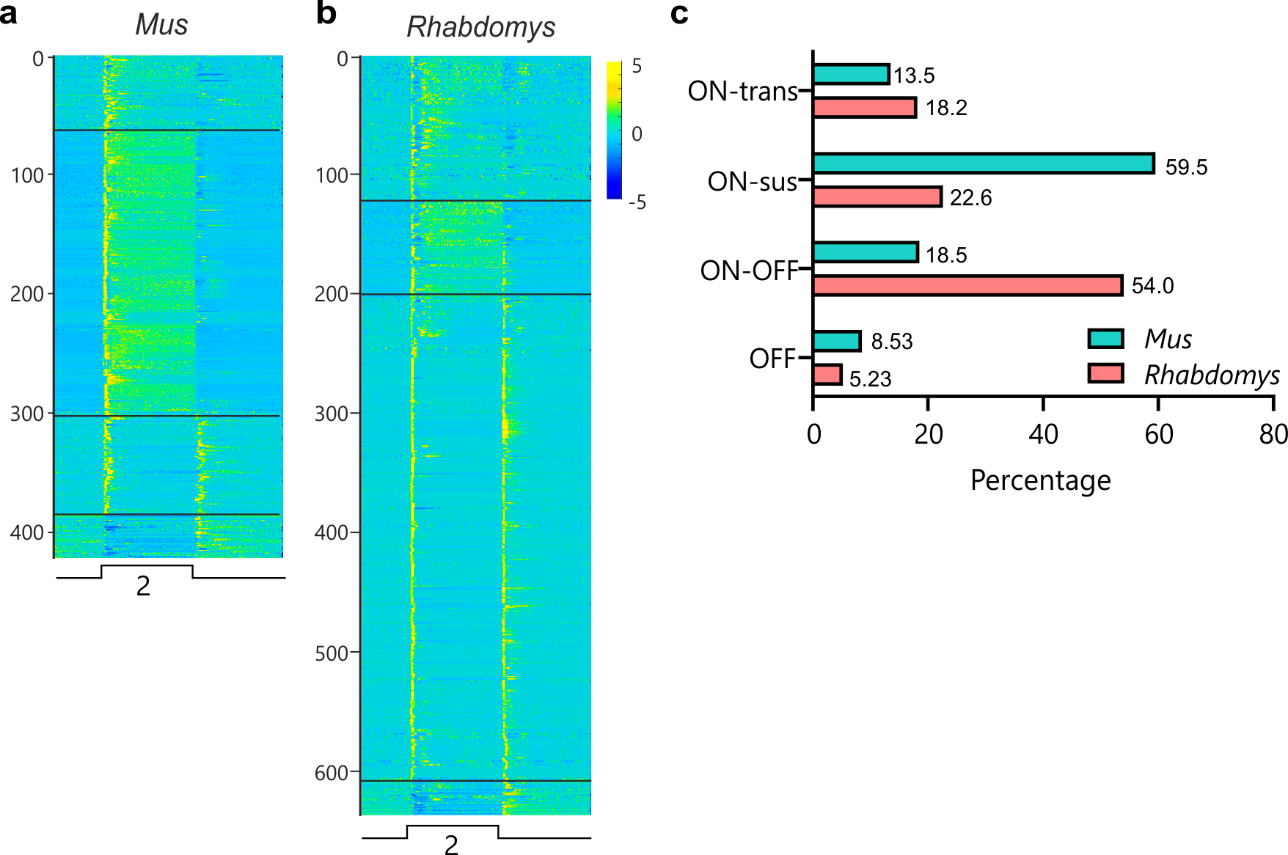

Supplementary Figure 3

**Responses to a 2s full field light step in *Mus* and *Rhabdomys*.**

**a&b)** Heat map showing mean light response in response to a 2s full-field step (97% Michelson contrast) for responsive neurons in *Mus* (a) and *Rhabdomys* (b), ordered as a function of response type (transient ON, sustained ON, on-off, off sustained). **c)** Pie charts showing proportion of units categorised into one of four response types: transient ON; Sustained ON; ON-OFF; OFF in *Mus* (right) and *Rhabdomys* (right).

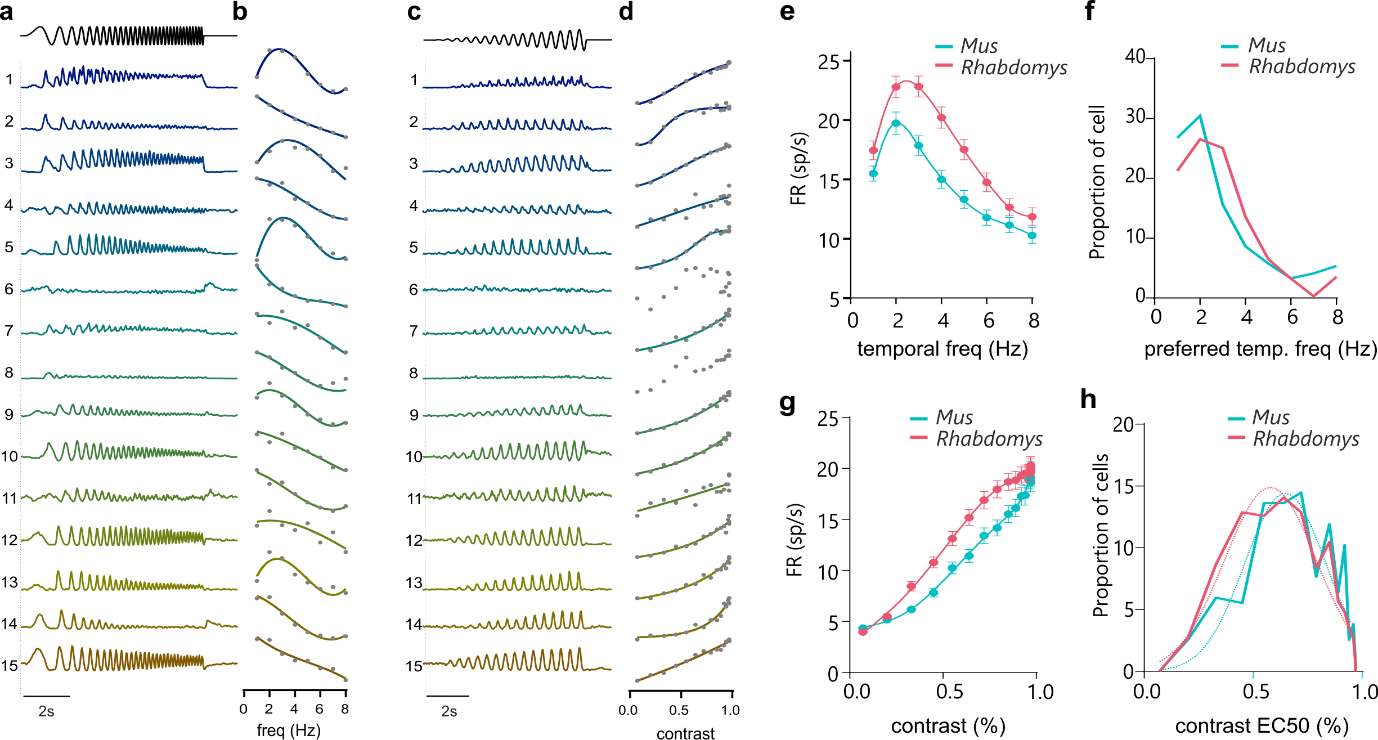

Supplementary Figure 4

**Temporal and contrast sensitivity of *Mus* and *Rhabdomys***

**a)** Mean response profile of clusters 1-15 to increasing frequency sinusoidal stimuli. **b)** Response amplitude to increasing frequency sinusoidal stimulus, binned at a resolution of 1Hz, for individual clusters. Data shows mean, normalised response profile. **c)** Mean response profile of clusters 1-15 to increasing contrast sinusoidal stimulus. **d)** Response amplitude to increasing contrast sinusoidal stimulus, for individual clusters. Data shows mean, normalised response profile. **e)** Response amplitude to increasing frequency sinusoidal stimulus, binned at a resolution of 1Hz, for *Mus* (cyan) and *Rhabdomys* (red). Data shows mean+/- SEM. **f)** Distribution of preferred temporal frequency for *Mus* (cyan) and *Rhabdomys* (red). **g)** Response amplitude to increasing contrast sinusoidal stimulus, shown for *Mus* (cyan) and *Rhabdomys* (red); data shows mean+/- SEM. **h)** Distribution of C50 values (half-maximum of Naka-Ruston curve) for *Mus* (cyan) and *Rhabdomys* (red).

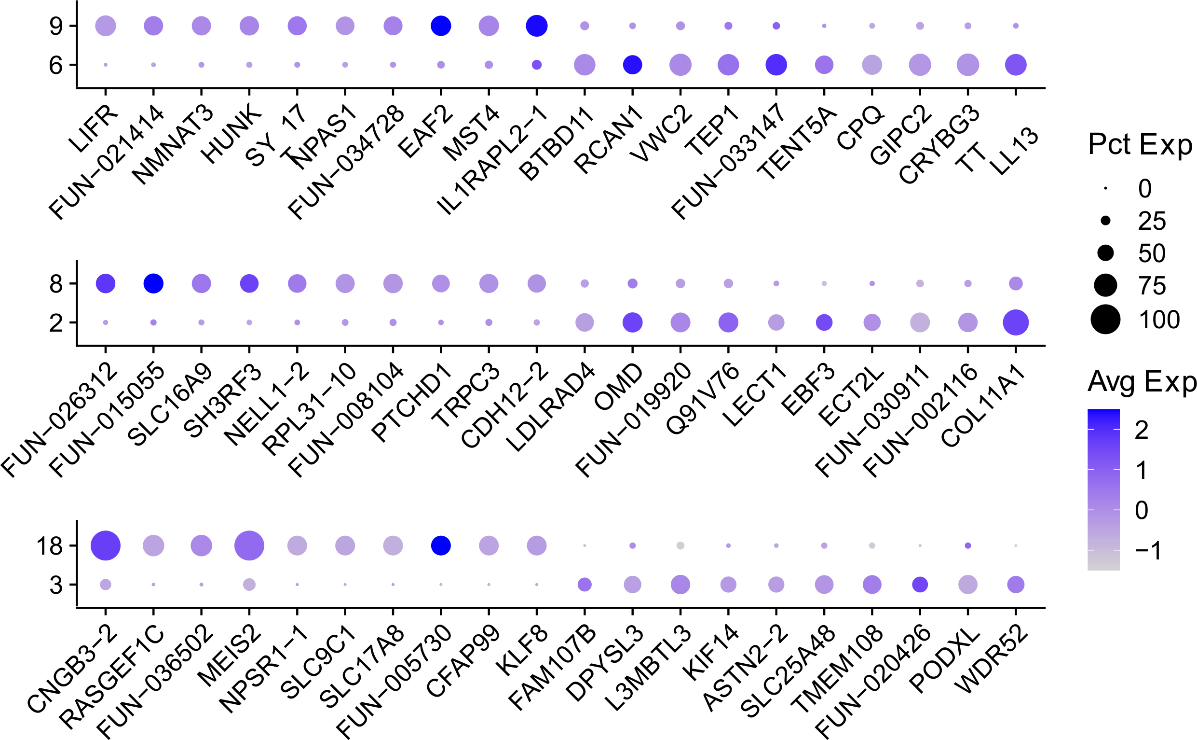

Supplementary Figure 5

Dotplots showing genes (columns) that distinguish pairs of *Rhabdomys* BC types (rows) that map to the same *Mus* BC type. The size of each circle represents the percentage of cells within a type that express the given gene, and the color represents the average expression of that gene within that cluster.

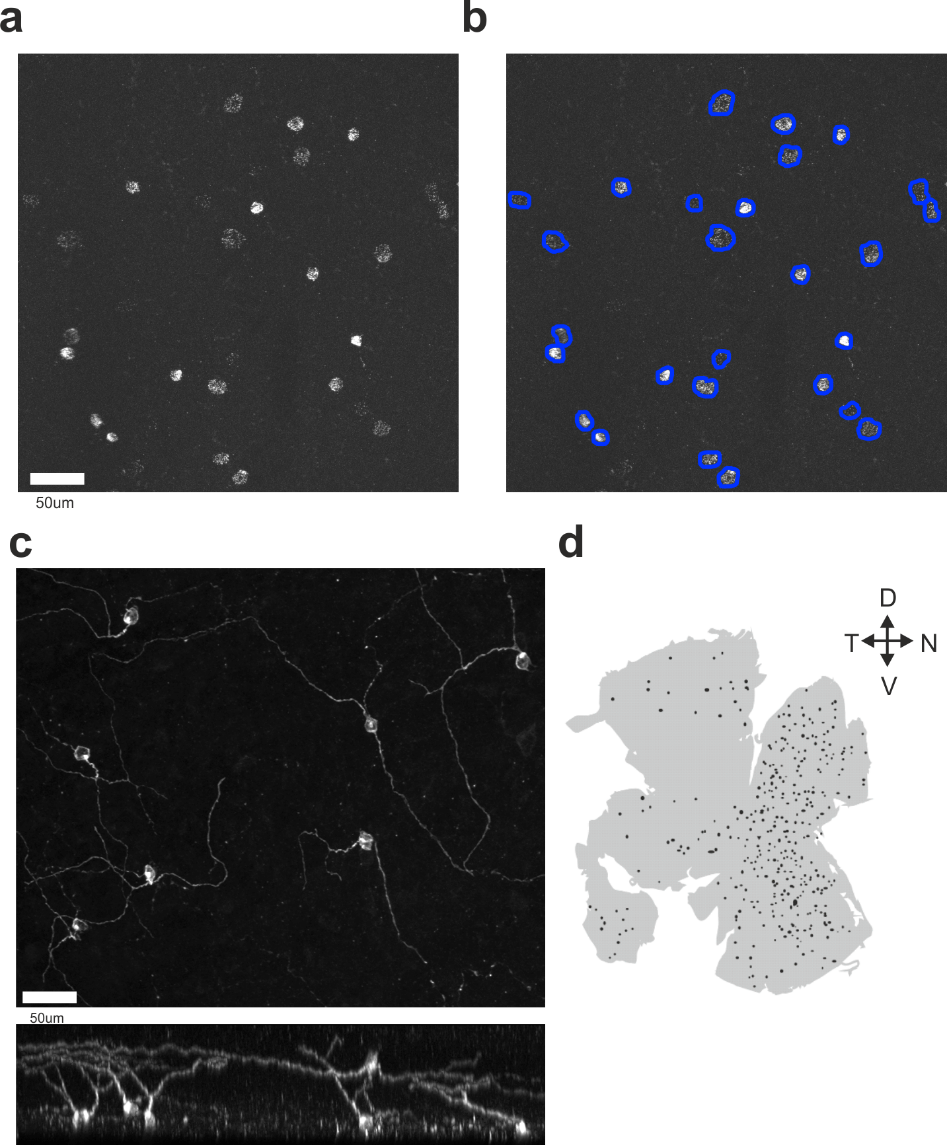

Supplementary Figure 6

**Automated cell identification**

**a)** Representative HCR-FISH in *Mus* retinal wholemount, with probe for *Mus Opn4*. Scale bar = 50µm. **b)** As in a), but with automated cell detection overlayed (blue edging). **c & d)** Immunohistochemistry for *Opn4* in *Rhabdomys* retina. High magnification (Scale bar = 50µm) (**c**) and representative retinal wholemount showing location of all Opn4 positive cell bodies in **d.**

| BC Type | *Rhab* 1 | *Rhab* 2 | *Mus* DropSeq | *Mus* ONC |
| --- | --- | --- | --- | --- |
| BC1A | 18.97 | 17.51 | 4.64 | 3.73 |
| BC1B | 2.61 | 1.83 | 3.37 | 1.40 |
| BC2 | 4.82 | 5.01 | 2.37 | 4.36 |
| BC3A | 8.17 | 9.31 | 2.28 | 5.31 |
| BC3B | 13.91 | 15.38 | 3.48 | 8.96 |
| BC4 | 6.41 | 6.53 | 1.69 | 3.38 |
| BC5A | 11.73 | 11.38 | 9.52 | 8.78 |
| BC5B | 4.42 | 3.98 | 2.04 | 5.63 |
| BC5C | 6.58 | 7.52 | 5.85 | 3.24 |
| BC5D | 2.24 | 1.92 | 2.35 | 2.30 |
| BC6 | 3.91 | 3.88 | 7.24 | 11.59 |
| BC7 | 8.17 | 7.92 | 7.49 | 5.33 |
| BC8 | 0.66 | 0.74 | 0.52 | 1.96 |
| BC9 | 2.12 | 2.21 | 0.80 | 2.03 |
| RBC | 5.27 | 4.87 | 46.35 | 31.97 |

**Supplementary Table 1. Bipolar cell frequencies across biological replicates of *Rhabdomys,* in comparison with *Mus.***

Rows show the percentage of *Rhabdomys* BCs mapping to each *Mus* BC type for two biological replicates*,* alongside previously published values for *Mus* BC classes ^1,2^.

| BC Type | *Rhab* 1 | *Rhab* 2 | *Mus* DropSeq | *Mus* ONC |
| --- | --- | --- | --- | --- |
| BC1A | 20.03 | 18.41 | 8.65 | 5.48 |
| BC1B | 2.76 | 1.93 | 6.28 | 2.06 |
| BC2 | 5.09 | 5.27 | 4.43 | 6.40 |
| BC3A | 8.63 | 9.79 | 4.24 | 7.81 |
| BC3B | 14.68 | 16.17 | 6.48 | 13.18 |
| BC4 | 6.77 | 6.87 | 3.16 | 4.97 |
| BC5A | 12.39 | 11.96 | 17.75 | 12.91 |
| BC5B | 4.66 | 4.19 | 3.80 | 8.28 |
| BC5C | 6.95 | 7.91 | 10.9 | 4.76 |
| BC5D | 2.36 | 2.02 | 4.39 | 3.39 |
| BC6 | 4.12 | 4.08 | 13.50 | 17.04 |
| BC7 | 8.63 | 8.33 | 13.95 | 7.83 |
| BC8 | 0.70 | 0.78 | 0.98 | 2.88 |
| BC9 | 2.24 | 2.33 | 1.49 | 2.99 |

**Supplementary Table 2. Cone bipolar cell frequencies across biological replicates of *Rhabdomys,* in comparison with *Mus.***

As in table 1, but with rod BCs excluded. Thus, rows show the percentage of *Rhabdomys* cone BCs mapping to each *Mus* cone BC type for two biological replicates*,* alongside previously published values for *Mus* BC classes ^1,2^.

| RGC Type | *Rhab* 1 | *Rhab* 2 | *Mus* 1 | *Mus* 2 | *Mus* 3 |
| --- | --- | --- | --- | --- | --- |
| C1_W3L1 | 8.00 | 10.33 | 8.81 | 8.91 | 7.58 |
| C2_W3D1 | 13.87 | 13.33 | 7.24 | 7.96 | 8.46 |
| C3_FminiON | 2.63 | 3.41 | 5.67 | 5.28 | 5.88 |
| C4_FminiOFF | 3.98 | 3.77 | 4.85 | 4.25 | 6.61 |
| C5_JRGC | 0.33 | 0.43 | 4.68 | 4.66 | 5.04 |
| C6_W3B | 7.47 | 7.18 | 3.80 | 4.92 | 4.24 |
| C7 | 1.19 | 1.52 | 4.44 | 3.78 | 5.19 |
| C8 | 2.11 | 2.76 | 2.98 | 3.57 | 3.76 |
| C9_TRGC_novel | 3.54 | 3.55 | 3.40 | 3.59 | 3.24 |
| C10 | 1.05 | 1.05 | 3.71 | 3.19 | 3.16 |
| C11 | 6.63 | 5.50 | 2.83 | 2.89 | 2.61 |
| C12_ooDSGC_N | 2.64 | 2.72 | 2.51 | 2.95 | 2.42 |
| C13_W3L2 | 3.65 | 3.29 | 2.62 | 2.48 | 2.85 |
| C14 | 0.87 | 0.88 | 2.64 | 2.60 | 2.17 |
| C15 | 1.83 | 1.69 | 2.39 | 2.57 | 2.27 |
| C16_ooDSGC_DV | 3.48 | 3.21 | 3.11 | 2.25 | 2.00 |
| C17_TRGC_S1 | 1.14 | 1.09 | 2.07 | 2.50 | 2.24 |
| C18 | 0.94 | 0.97 | 2.11 | 2.79 | 1.84 |
| C19 | 4.44 | 4.35 | 2.01 | 1.90 | 2.58 |
| C20 | 1.99 | 1.47 | 1.80 | 2.00 | 2.08 |
| C21_TRGC_S2 | 4.46 | 4.72 | 2.39 | 1.37 | 2.35 |
| C22_MX | 0.24 | 0.33 | 1.19 | 1.90 | 1.74 |
| C23_W3D2 | 4.11 | 3.37 | 2.01 | 1.22 | 2.07 |
| C24 | 3.93 | 3.53 | 2.05 | 1.58 | 1.25 |
| C25 | 2.07 | 1.81 | 1.85 | 1.75 | 1.07 |
| C26 | 2.28 | 1.83 | 1.73 | 1.61 | 1.23 |
| C27 | 1.17 | 1.07 | 1.72 | 1.57 | 1.25 |
| C28_FmidiOFF | 0.49 | 0.54 | 1.20 | 1.63 | 1.36 |
| C29 | 2.64 | 2.73 | 1.35 | 1.63 | 1.15 |
| C30_W3D3 | 1.42 | 1.83 | 1.45 | 1.03 | 1.75 |
| C31_M2 | 0.18 | 0.12 | 0.91 | 1.58 | 1.01 |
| C32_FRGC_novel | 0.53 | 0.58 | 1.51 | 1.11 | 0.98 |
| C33_M1a | 0.03 | 0.08 | 0.79 | 0.91 | 0.96 |
| C34 | 1.14 | 1.01 | 1.04 | 0.63 | 1.08 |
| C35 | 1.32 | 1.50 | 0.86 | 0.97 | 0.75 |
| C36 | 0.21 | 0.12 | 0.81 | 0.64 | 0.61 |
| C37 | 0.23 | 0.24 | 0.85 | 0.67 | 0.38 |
| C38_FmidiON | 0.17 | 0.24 | 0.56 | 0.65 | 0.50 |
| C39 | 1.00 | 1.06 | 0.57 | 0.37 | 0.80 |
| C40_M1b | 0.02 | 0.01 | 0.26 | 0.56 | 0.52 |
| C41_AlphaONT | 0.06 | 0.15 | 0.31 | 0.42 | 0.30 |
| C42_AlphaOFFS | 0.31 | 0.24 | 0.41 | 0.34 | 0.24 |
| C43_AlphaONS_M4 | 0.08 | 0.12 | 0.25 | 0.42 | 0.18 |
| C44 | 0.10 | 0.21 | 0.10 | 0.20 | 0.16 |
| C45_AlphaOFFT | 0.03 | 0.07 | 0.16 | 0.20 | 0.08 |

**Supplementary Table 3. Retinal ganglion cell frequencies across biological replicates of *Rhabdomys,* in comparison with *Mus***

Each row shows the percentage of RGCs mapping to each *Mus* RGC type for two biological replicates of *Rhabdomys,* and three biological replicates of *Mus* ^2^.

**References**

1	Shekhar, K. *et al.* Comprehensive Classification of Retinal Bipolar Neurons by Single-Cell Transcriptomics. *Cell* **166**, 1308-1323 e1330, doi:10.1016/j.cell.2016.07.054 (2016).

2	Tran, N. M. *et al.* Single-Cell Profiles of Retinal Ganglion Cells Differing in Resilience to Injury Reveal Neuroprotective Genes. *Neuron* **104**, 1039-1055 e1012, doi:10.1016/j.neuron.2019.11.006 (2019).
